# BK channel gain-of-function disrupts limb control by suppressing neurotransmission during a critical developmental window

**DOI:** 10.1101/2023.09.20.558625

**Authors:** Simon A. Lowe, Abigail D. Wilson, Gabriel Aughey, Animesh Banarjee, Talya Goble, Nell Simon-Batsford, Angelina Sanderson, Patrick Kratschmer, Maryam Balogun, Hao Gao, Sherry S. Aw, James E.C. Jepson

## Abstract

Gain-of-function mutations in BK potassium channels (BK GOF) cause debilitating involuntary limb movements. BK channels modulate action potential shape and neurotransmission in mature neurons, yet some BK GOF mutations also cause neurodevelopmental morbidities. Thus, whether BK GOF impairs limb control by altering the excitation/inhibition of mature motor circuits, or by disrupting their development, remains unclear. To address this issue, we developed a genetic method enabling spatiotemporal control of BK channel expression in neurons of the fruit fly, *Drosophila*. In concert with high-resolution measurements of limb kinematics, we demonstrate that GOF BK channels act during a narrow neurodevelopmental period to perturb limb control in adult flies. During this period, BK GOF alters synaptic localisation of the key active zone protein Bruchpilot and suppresses excitatory neurotransmission. In a wild-type background, we find that reducing neural activity during neurodevelopment yields similar motor defects to those observed in BK GOF flies. Conversely, enhancing neural excitability during development rescues alterations in limb kinematics in BK GOF flies. Collectively, our results suggest that BK GOF perturbs limb control largely by disrupting activity-dependent aspects of neuronal development.

## Introduction

BK (Big K^+^) channels are a highly conserved member of the voltage-gated potassium channel (VGPC) gene family ^1^. So-called for their strikingly high single-channel conductance, BK channels are unique amongst VGPCs in their ability to couple increases in intracellular Ca^2+^ to K^+^ efflux across membranes ^2^, a feat accomplished by two C-terminal regulator of K^+^ conductance domains (RCK1 and RCK2) that bind Ca^2+^ and facilitate conformational changes that enhance open channel probability ^3^.

BK channels are broadly expressed throughout Bilaterian nervous systems ^4–6^ and their activation has complex and often cell-specific effects on neuronal physiology ^2^. For example, axonal BK channels facilitate action potential (AP) repolarisation and gate a fast afterhyperpolarisation (fAHP) current that influences AP frequency ^7^, while presynaptic BK channels suppress neurotransmitter release by inactivating synaptic voltage-gated Ca^2+^ channels (VGCCs) ^8^. BK channel expression and function can also be modulated by neuronal activity ^9,10^. BK channels thus modulate numerous aspects of neural coding, including spike frequency, synaptic strength, and homeostatic alterations in intrinsic excitability ^8–11^.

A growing number of studies have shown that enhancement of BK channel activity via inherited or de novo gain-of-function (GOF) mutations causes dystonia and dyskinesia in humans ^12–14^, movement disorders characterised by sustained or transient involuntary muscle contractions and a debilitating loss of postural control ^15,16^. However, the mechanisms by which BK channel GOF mutations disrupt motor control remain poorly understood. The most-studied GOF variant is a dominant aspartic acid to glycine change at residue 434 (D434G), located within the BK channel α-subunit RCK1 domain ^12^. D434G enhances BK channel activity by increasing the Ca^2+^ sensitivity of the channel ^17,18^, and heterozygosity for D434G in humans results in a complex syndrome characterised by bouts of dystonic and dyskinetic movements in the limbs, mouth, and tongue ^12^. Recent work in corresponding mouse models has also shown that D434G heterozygosity enhances the firing rate of murine cerebellar Purkinje neurons and dentate granule cells by accelerating the kinetics of the fAHP ^19,20^.

These data suggest that GOF BK channels may perturb motor control by altering the excitation/inhibition balance of pre-motor brain regions such as the cerebellum. However, other BK channel GOF mutations in humans are associated with developmental delay and intellectual disability as well as involuntary movements ^13,21^, raising the possibility that BK channel GOF could instead cause motor defects by affecting neuronal development. Indeed, BK channels are expressed in the developing post-natal murine brain ^22,23^, and during this period, stimulus-independent Ca^2+^ transients that drive activity-dependent aspects of neuronal growth occur widely and are capable of recruiting BK channels ^24^.

We therefore sought to disentangle the relative effects on movement and limb control of enhancing BK channel activity in the developing versus the mature nervous system. *Drosophila* represents an ideal model to achieve this goal due to its wealth of well-established methods to control gene expression in time and space ^25^. Furthermore, we recently generated a *Drosophila* knock-in model of the D434G mutation and demonstrated that flies heterozygous for the equivalent amino-acid change (E366G) in the SLOWPOKE (SLO) BK channel α-subunit (*slo*^E366G/+^) exhibit an array of motor defects, including reduced total movement, reduced locomotor speed, and bouts of dyskinesia-like leg twitches ^26^. In this background, we developed a genetic strategy that allowed us reversibly promote or inhibit expression of neuronal GOF BK channels. Combining this method with measurements of limb kinematics, synaptic protein expression, and neurotransmitter release, we find that GOF BK channels predominantly act during the final stages of neurodevelopment to disrupt limb control in adult flies, and do so in part by suppressing neural activity during this stage.

## Results

### Reversible control of BK channel expression in *Drosophila* neurons

We developed a method to reversibly tune neuronal expression of SLO BK channels through post-translational control, while leaving *slo* mRNA expression under control of its endogenous cis-regulatory elements. We based our approach on our previous studies of *dyschronic* (*dysc*). *dysc* encodes a scaffold protein that binds to SLO BK channels and promotes their expression; brain-wide SLO expression is hence greatly reduced in flies homozygous for a *dysc* LOF allele (*dysc*^s168^) ^27^. Unlike *slo* null flies ^28^, *dysc*^s168^ homozygotes do not exhibit overt motor defects ^26^, likely because SLO expression is not completely abolished in *dysc*^s168/s168^ flies ^27^. Indeed, we recently showed that homozygosity for *dysc*^s168^ suppressed motor defects in *slo*^E366G/+^ flies by reducing the expression of GOF BK channels^26^. Furthermore, restoring DYSC expression solely in post-mitotic neurons (*nsyb*-Gal4 > UAS-*dysc*) restored locomotor defects to *slo*^E366G/+^, *dysc*^s168/s168^ flies ^26^, showing that GOF BK channels act in post-mitotic neurons to disrupt movement.

In this background we placed UAS-*dysc* under thermogenetic control using *tub*-Gal80^ts^ ^29^. This construct encodes GAL80^ts^, a globally expressed GAL4-inhibitor that blocks GAL4-driven transgene expression at low (18°C) but not high temperatures (29°C), and which has previously been used to limit transgene expression to the larval ^30^, pupal ^31^, or adult ^32,33^ stages of the *Drosophila* lifecycle. This method should allow us to promote robust expression of DYSC, and therefore SLO BK channels, at the permissive temperature of 29°C and suppress expression at 18°C (Fig. 1a-b; see Materials and Methods for further notes). Using an antibody against human hSlo1 BK channels that also binds to *Drosophila* SLO, we verified GAL80^ts^ enabled specific enhancement of neuronal BK channel expression at 29°C relative to 18°C at both the pupal and adult stages (Supplementary Fig. S1a-g).

**Fig. 1.**
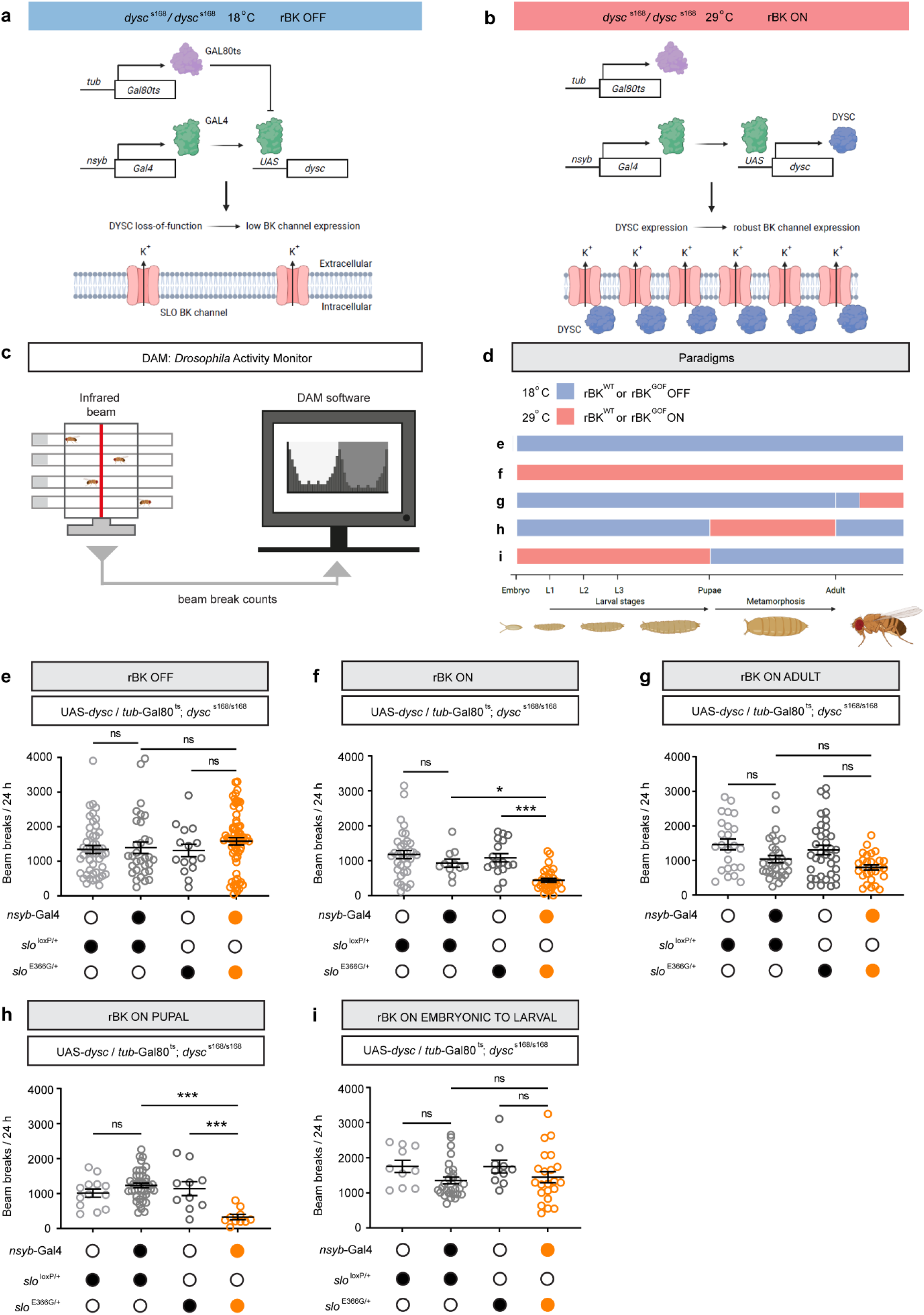
Expression of GOF BK channels during neurodevelopment reduces spontaneous locomotion. **(a-b)** Schematic illustrating a genetic method enabling temperature-dependent control of BK channel expression. **(c)** Schematic illustrating the DAM system and representative data. Flies are loaded individually into behavioural tubes. The DAM system records the number of times an individual fly breaks an infrared beam over 24 h in a 12 h light: 12 h dark cycle. **(d)** Schematic illustrating developmental stages during which neuronal rBK^WT^ or rBK^GOF^ expression is induced. Letters refer to data shown in subsequent panels e-i. Experiments on adult male flies took place 5-7 days after eclosure. **(e-i)** Total number of beam breaks in a 24 h period, recorded using the DAM system, in the paradigms illustrated in (d). Experimental genotype is denoted in orange, control genotypes in grey. Developmental stages during which rBK expression is induced are noted above each dataset. Error bars: SEM. *p< 0.05, *** p< 0.0005, ns – p> 0.05, Kruskal-Wallis non-parametric test with Dunn’s multiple comparisons (e, f, h, i) or one-way ANOVA with Tukey’s multiple comparisons test (g). n-values are: (e) n = (left to right) 46, 32, 15, 69 flies; (f) n = 35, 11, 17, 32 flies; (g) n = 24, 33, 36, 27 flies; (h) n = 13, 40, 10, 10 flies; (i) n = 10, 28, 10, 22 flies.

For simplicity, we will henceforth use the terms ‘rBK ON/OFF’ (robust BK channel expression on or off) to denote the temperature-dependent promotion or suppression of SLO BK channel expression. This will be performed in backgrounds heterozygous for either the BK GOF allele (*slo*^E366G/+^: rBK^GOF^) or a genetically isogenised wild-type *slo* allele derived using the same homologous recombination process (*slo*^loxP/+^: rBK^WT^) ^26^.

### GOF BK channels act during development to disrupt movement in *Drosophila*

To confirm our method yielded expected behavioural outcomes, we tested whether constitutively inhibiting neuronal rBK^GOF^ expression throughout the *Drosophila* lifecycle suppressed locomotor defects of *slo*^E366G/+^ flies ^26^. By quantifying the number of infrared beam breaks made by flies housed in *Drosophila* Activity Monitor (DAM) systems ^34^ over 24 h (Fig. 1c), we indeed found a robust suppression of motor defects in *slo*^E366G/+^ flies using this approach (Fig. 1d, e). Conversely, constitutively promoting neuronal rBK^GOF^ expression restored locomotor defects in this background, as shown by a significant reduction in beams breaks compared to relevant controls (Fig. 1d, f).

We then tested whether inducing rBK^GOF^ expression specifically in the mature adult nervous system was sufficient to cause locomotor defects. Since synaptic growth during neurodevelopment extends into the first 1-2 days of fruit fly adulthood ^35^, we induced rBK^GOF^ expression after 2 days of adulthood and maintained expression for 3 days prior to locomotor analysis, allowing an extended period for GOF BK channels to accumulate and acutely induce motor deficits (Fig. 1d; we henceforth define this condition as ‘adult neurons’). Surprisingly, we found no difference in overall movement between flies with rBK^GOF^ or rBK^WT^ expression induced in adult neurons (Fig. 1g). We therefore moved backwards through developmental time, inducing neuronal rBK^GOF^ expression solely during either pupal stage of the fly lifecycle, or the embryonic through larval stages (Fig. 1d). We found that inducing neuronal rBK^GOF^ (but not rBK^WT^) expression during the pupal stage led to striking locomotor defects in resulting adult flies (Fig. 1h), whereas inducing neuronal rBK^GOF^ expression during the embryonic to larval stages of the fly life cycle had no effect (Fig. 1i).

Measurements from the DAM system were obtained across a complete day-night cycle. Hence, alterations in the pattern of locomotor activity during day/night periods, rather than locomotor capacity per se, could contribute to the reduced movement caused by induction of rBK^GOF^ expression during the pupal stage. To rule out this possibility, we deployed a video-based system called DART (*Drosophila* Arousal Tracking) capable of continuously monitoring fly movement before and after an induced mechanosensory stimulus ^36^ (Fig. 2a). Using DART, we previously showed that *slo*^E366G/+^ flies exhibit significantly less movement in response to a startle-inducing mechanosensory stimulus ^26^. Interestingly, we found no difference in the stimulus-induced peak locomotor velocity of flies with rBK^GOF^ or rBK^WT^ expression induced in adult neurons (Fig. 2b-d). In contrast, inducing neuronal rBK^GOF^ expression during the pupal stage strongly reduced locomotor velocity compared to rBK^WT^ controls (Fig. 2b, e-f). Thus, GOF BK channel expression solely during the pupal stage is sufficient to disrupt both self-driven and stimulus-induced movement.

**Fig.2. Expression.**
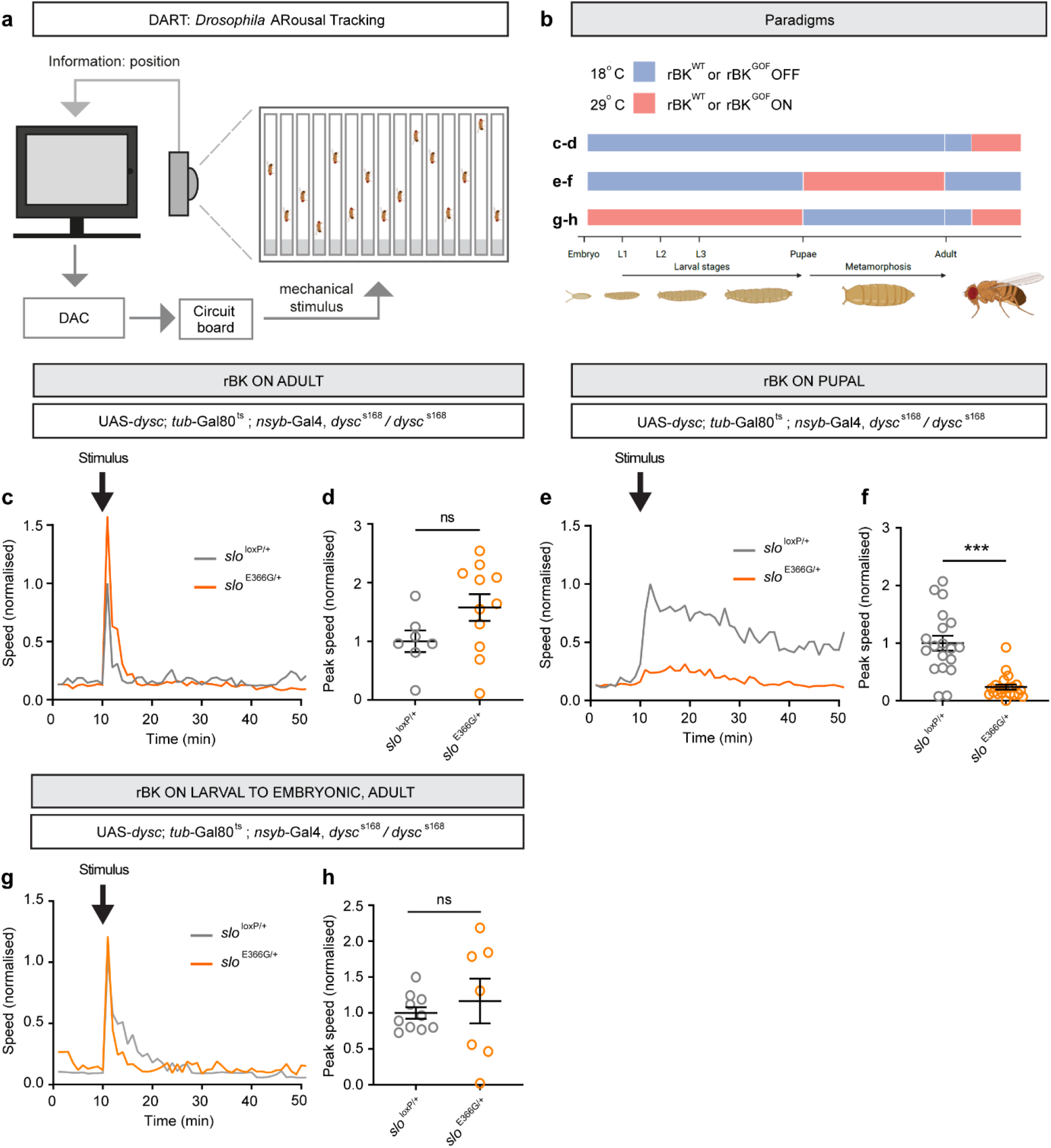
of GOF BK channels during neurodevelopment reduces stimulus-induced locomotion. **(a)** Schematic illustrating a stimulus-dependent locomotor assay: the *Drosophila* ARousal Tracking (DART) system. **(b)** Schematic illustrating developmental stages during which neuronal rBK^WT^ or rBK^GOF^ expression is induced. Letters refer to data shown in panels C-H. Experiments on adult male flies took place 5-7 days after eclosure. **(c, e, g)** Traces showing average speed over time in one-minute bins (mm/s) for control (n = 7) and *slo*^E366G/+^ (n = 11) flies in the paradigm shown in (b). Mechanical stimulus was delivered at 10 minutes. **(d, f, h)** Quantification of speed following mechanical stimulus. Each data point represents average speed (mm / s) in the 1 min period immediately following stimulation for each fly. Data are presented normalised to the control mean. n = 7, 11 flies. Error bars: SEM. n-values: c-d: n = 7 *slo*^loxP/+^, 11 *slo*^E366G/+^; e-f: n = 19, 21; g-h: n = 10, 7. ***p < 0.0005, ns – p > 0.05, t-test with Welch’s correction (d) or Mann-Whitney U-test (f, h).

To test whether expression of GOF BK channels specifically during the pupal stage was necessary as well as sufficient to recapitulate the effect of constitute expression, we induced rBK^GOF^ expression in all stages of the fly life cycle except the pupal stage and quantified the impact on self-driven and stimulus-induced locomotion using the DAM and DART systems respectively. Using the DAM system, we observed a significant reduction in overall movement in flies with neural rBK^GOF^ expression induced during the embryonic-to-larval, and adult stages alone (Supplementary Fig. S2a, b). However, this phenotype was far less severe compared to the converse experiment of inducing rBK^GOF^ expression solely in the pupal stage, as quantified by Cohen’s effect size *d* (embryonic-to-larval and adult stage induction: *d* = -0.59; pupal induction: *d* = -2.62). Furthermore, while inducing rBK^GOF^ expression in embryonic-to-larval and adult neurons appeared to increase the variability of stimulus-induced locomotor velocity (as measured by DART), mean velocity was not reduced compared to rBK^WT^ controls (Fig. 2g, h), in contrast to the effect of inducing neuronal rBK^GOF^ expression during the pupal stage (Fig. 2e, f). These data suggest that GOF BK channel expression during a critical period prior to adulthood – during the development of the adult nervous system – is responsible for the gross disruptions in locomotor capacity in adult flies.

### GOF BK channels act during the late stages of neurodevelopment to perturb movement

The transformation of the larval into the adult *Drosophila* nervous system during metamorphosis occurs via stereotyped, temporally organised phases of neuronal development, which include the pruning of existing larval dendrites and axons; the re-growth of dendritic/axonal arbors into their adult forms; and finally, the formation and maturation of synaptic connections ^37,38^. We searched for clues as to which of these processes BK GOF might perturb.

To do so, we induced neuronal rBK^GOF^ expression during five 24 h windows coinciding with distinct neurodevelopmental transitions during the pupal stage (Fig. 3a-b; note that under these thermal conditions, the pupal stage takes eight days to complete). We observed a clear time-dependent effect of neuronal rBK^GOF^ expression during the pupal stage on subsequent adult fly movement (Fig. 3b). Induction of neuronal rBK^GOF^ expression during early-to-mid stages of the pupal stage either did not significantly reduce movement compared to controls lacking rBK^GOF^ expression (early stage), or reduced movement only to a small degree (mid-stage) (Fig. 3b). In contrast, induction of neuronal rBK^GOF^ expression in neurons for 24 h during either of the last two days (day 6-7 and day 7-8) of the pupal stage profoundly reduced movement in resulting adult flies (Fig. 3b). Similarly inducing neuronal rBK^WT^ expression for 24 h during the late pupal stage, in contrast, had no effect on overall movement (Supplementary Fig. S2c, d). We also confirmed the detrimental effect of inducing neuronal rBK^GOF^ expression during the late pupal stage on movement using DART (Supplementary Fig. S3a-c). These data demonstrate that GOF BK channels act during a restricted neurodevelopmental window within late metamorphosis to impact movement.

**Fig. 3.**
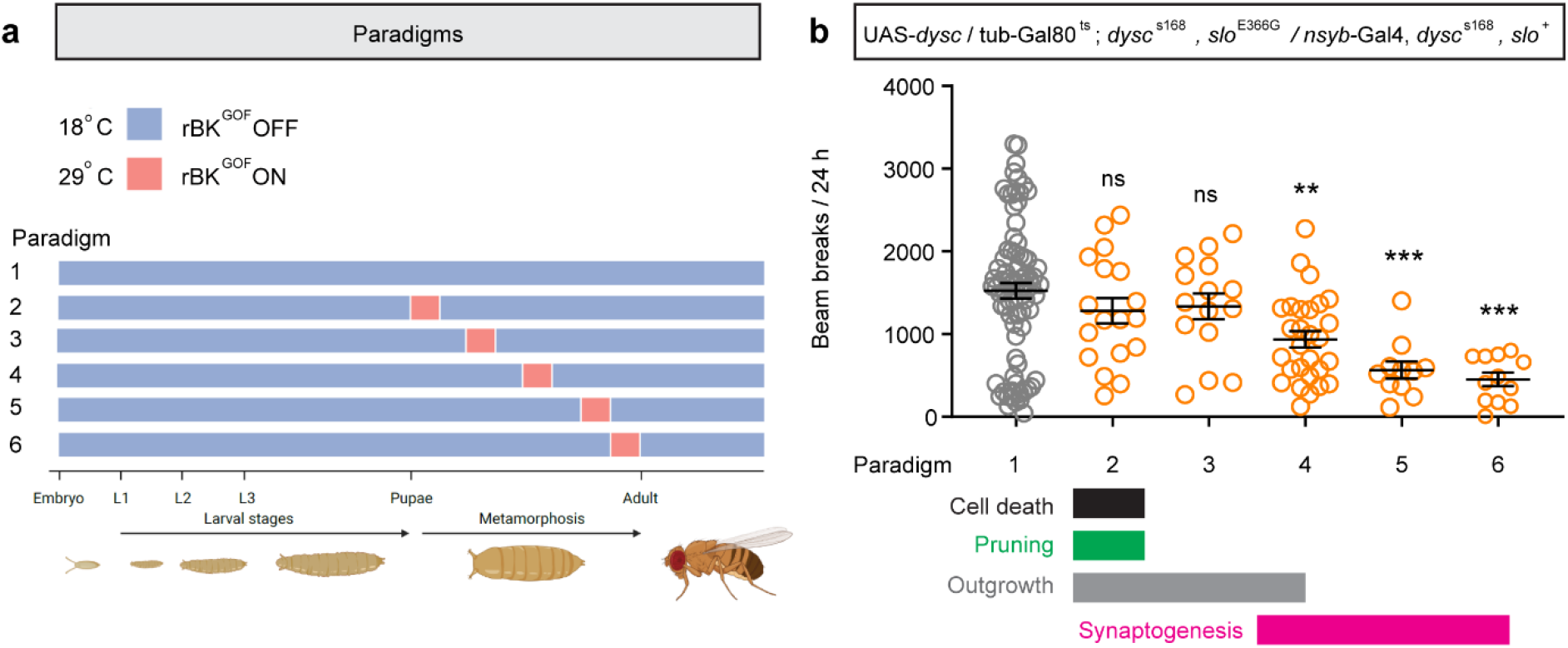
Neuronal expression of GOF BK channels during late development decreases spontaneous locomotion. **(a)** Schematic illustrating induction of neuronal rBK^GOF^ expression for 24 h periods during distinct periods of the pupal stage (paradigms 2-6; paradigm 1 denotes continued suppression of rBK^GOF^ expression). Experiments on adult male flies took place 5-7 days after eclosure. **(b)** Total number of beam breaks in a 24 h period recorded using the DAM system in the paradigms illustrated in (a). n = (left to right) 81, 18, 15, 28, 11, 12 flies. Below graph: approximate timeline showing developmental processes overlapping with the period of GOF BK channel expression in each paradigm. Error bars: SEM. **p< 0.005, *** p< 0.0005, ns – p> 0.05, Kruskal-Wallis non-parametric test with Dunn’s multiple comparisons.

### Enhancing BK channel activity during the late pupal stage disrupts limb kinematics

We next sought to understand how enhancing BK channel activity during neurodevelopment impacted movement at the level of individual limbs. Human carriers of the equivalent mutation to *slo*^E366G/+^ (hSlo1^D434G/+^) present with paroxysmal non-kinesigenic dyskinesia (PNKD), a movement disorder characterised by bouts of involuntary dystonic and choreiform limb movements ^12^. *slo*^E366G/+^ flies exhibit similar bouts of leg twitches, consisting of rapid, repetitive movements isolated to a single limb – a phenotype not observed in control flies ^26^. We therefore asked whether inducing GOF BK channel expression during neurodevelopment also yielded dyskinesia-like phenotypes. Indeed, inducing neuronal rBK^GOF^ expression during the late pupal stage caused at least one bout consisting of multiple leg twitches during a 5 minute recording period in 66% of flies examined (n = 9), whereas inducing rBK^GOF^ expression in adult neurons did not (n = 5; Supplementary Fig. S3d-g).

Dyskinesia-like phenotypes in the above backgrounds occurred while flies were at rest. Hence, this analysis did not reveal how alterations in limb kinematics contributed to the reductions in locomotor velocity observed in *slo*^E366G/+^ flies ^26^ and in flies with neuronal rBK^GOF^ expression induced during the pupal stage (Fig. 2e, f). To study in detail how BK GOF during neurodevelopment impacted limb movement we utilised FLLIT (Feature Learning-based LImb segmentation and Tracking), a machine-learning image analysis package capable of deriving gait parameters from high-speed videography of fly locomotion ^39^ (Fig. 4a; see Materials and Methods and refs. ^39,40^ for descriptions of gait parameters). Using FLLIT, we first examined limb movements in *slo*^E366G/+^ flies versus controls (Fig. 4b, c). We uncovered changes in four kinematic parameters that likely contribute to the reduced movement and locomotor speed observed in *slo*^E366G/+^ flies ^26^. These are: a reduced percentage of time each limb spent moving (Fig. 4d); reduced stride length (displacement; Fig. 4e); reduced velocity of limb movement (Fig. 4f); and an enhanced non-linearity of limb movement (calculated by normalising the actual limb path to the idealised limb displacement) suggestive of uncoordinated limb movements (Fig. 4b-c, g).

**Fig. 4.**
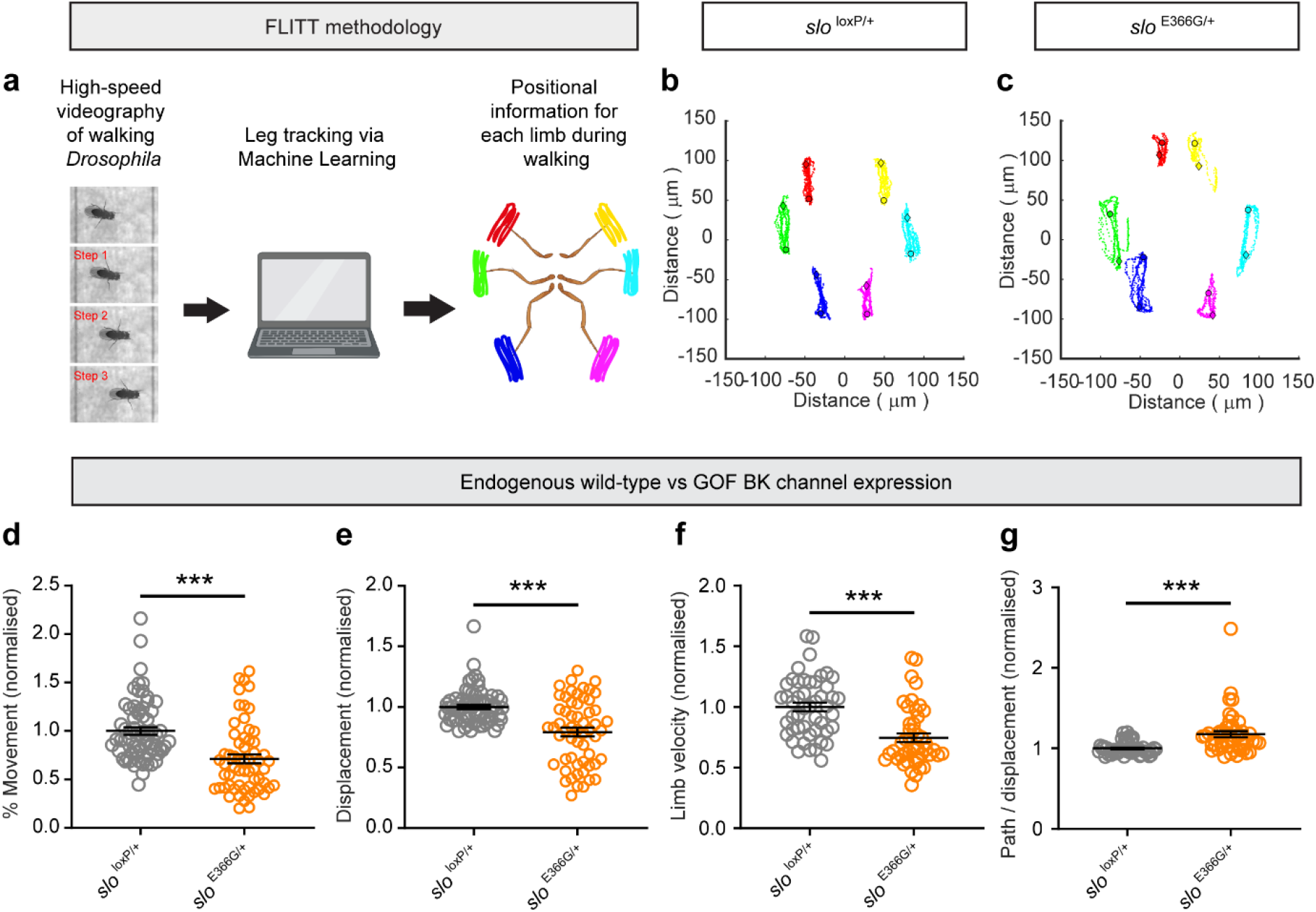
Flies expressing GOF BK channels exhibit alterations in limb kinematics. **(a)** Schematic illustrating the FLLIT methodology. Schematic on right shows overlaid movement of each limb over multiple strides, relative to the centre of the body. **(b, c)** Representative FLLIT-derived traces showing movement of each limb over 3-6 strides, relative to the centre of the body, for control (*slo*^loxP/+^) (b) and BK GOF (*slo*^E366G/+^) (c) adult male flies. **(d-g)** FLLIT-derived gait parameters comparing *slo*^loxP/+^ and *slo*^E366G/+^ flies. See Materials and Methods for detailed descriptions of gait parameters. Data points represent mean values from a single limb across 3-6 strides. Data are presented normalised to control means. n = 66 limbs across 11 flies (*slo*^loxP/+^) and 60 limbs across 10 flies (*slo*^E366G/+^). Error bars: SEM. *p<0.05, *** p< 0.0005, ns – p> 0.05, Mann-Whitney U-test.

We next compared the effect of inducing neuronal rBK^GOF^ expression in adult neurons or during late neurodevelopment on limb kinematics. rBK^GOF^ expression in adult neurons caused a small, marginally significant reduction in the time each limb spent moving (Fig. 5a-d), but no change in stride displacement, limb velocity, or the non-linearity of limb movement (Fig. 5e-g). In contrast, induction of neuronal rBK^GOF^ expression during late neurodevelopment significantly reduced the time each limb spent moving, the stride displacement, and limb velocity; and significantly enhanced the non-linearity of limb movement (Fig. 5h-n).

**Fig. 5.**
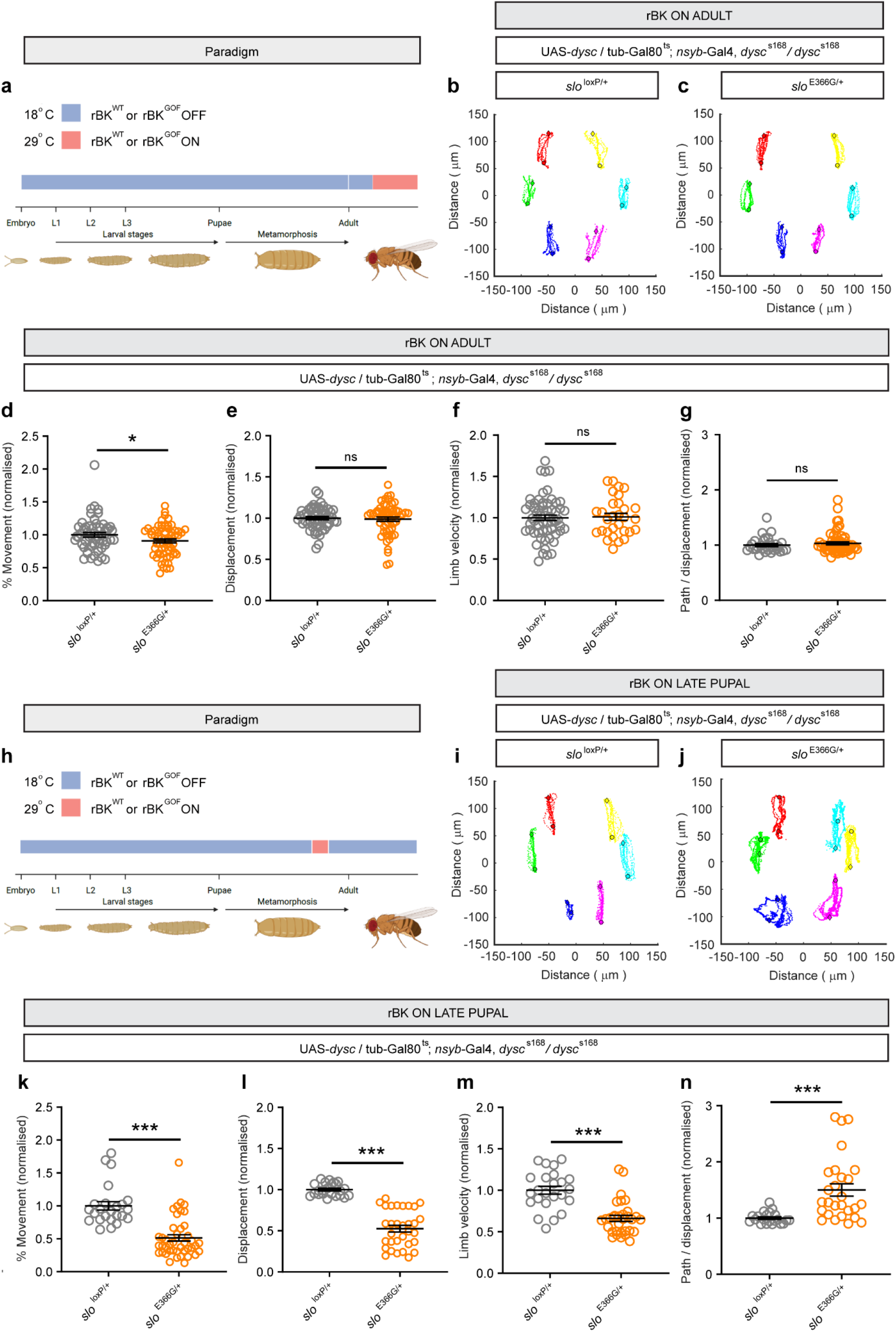
Neuronal expression of GOF BK channels during late development perturbs limb kinematics. **(a)** Schematic illustrating paradigm to induce rBK^WT^ or rBK^GOF^ expression in adult neurons. Experiments on adult male flies took place 5-7 days after eclosure. **(b, c)** Representative FLLIT-derived traces showing movement of each limb over 3-6 strides, relative to the centre of the body, for flies with rBK^WT^ (b) or rBK^GOF^ (c) expression induced in adult neurons. **(d-g)** FLLIT-derived gait parameters for flies with robust expression of neuronal wild type or GOF BK channels restricted to the adult stage. n = 54 limbs across 9 flies (control, grey) and 60 limbs across 10 flies (BK GOF, orange). **(h)** Schematic illustrating paradigm to induce neuronal rBK^WT^ or rBK^GOF^ expression during late neurodevelopment. Experiments on adult male flies took place 5-7 days after eclosure. **(i, j)** Representative FLLIT-derived traces showing movement of each limb over 3-6 strides, relative to the centre of the body, for flies with neuronal rBK^WT^ or rBK^GOF^ expression induced during a 24 h period at the end of the pupal stage. **(k-n)** FLLIT-derived gait parameters for flies with neuronal rBK^WT^ or rBK^GOF^ expression induced during a 24 h period at the end of the pupal stage. n = 24 limbs across 4 flies (control, grey) and 42 limbs across 7 flies (BK GOF, orange). Error bars: SEM. *p<0.05, *** p< 0.0005, ns – p> 0.05, Mann-Whitney U-test.

We also noted a substantial increase in the variability of limb displacement and movement linearity in *slo*^E366G/+^ flies, quantified via the coefficient of variation (Supplementary Fig. S4a, b). A small increase in the variability of limb movements was observed following induction of rBK^GOF^ in adult neurons (Supplementary Fig. S4c-e), while a much greater increase was observed following induction of neuronal rBK^GOF^ expression during late neurodevelopment (Supplementary Fig. S4f-g).

Collectively, these data demonstrate that inducing neuronal rBK^GOF^ expression solely during the late pupal stage causes dyskinesia-like limb movements and recapitulates the phenotype observed following constitutive GOF BK channel expression, while expression in adult neurons does not.

### BK channel GOF alters presynaptic protein expression during the late pupal stage

The critical neurodevelopmental window in which GOF BK channels act to perturb adult movement coincides with periods of neural arborisation, synaptogenesis, and synaptic maturation, within the pupal stage (Fig. 3b) ^37,38^, suggesting that one or more of these processes may be disrupted by BK channel GOF. To test this, we first examined axonal and synaptic development in flies with constitutive expression of GOF BK channels (*slo*^E366G/+^) compared to controls (*slo*^loxP/+^). We focused our analysis on two neural subtypes in the fly brain – the mushroom bodies (MBs) and PDF-positive large ventral lateral neurons (l-LN_v_s) – since SLO BK channels can be visualised in the MBs ^27^ and influence action potential repolarisation in the l-LN_v_s ^26,41^. We found no difference in axonal growth of four mushroom body compartments (the αβ and α’β’ lobes) between *slo*^E366G/+^ and control adult flies (Supplementary Fig. S5a-f). Presynaptic boutons from the l-LN_v_s are broadly dispersed throughout the fly optic lobe, allowing us to quantify their number and size. Neither of these synaptic parameters were altered between *slo*^E366G/+^ and control flies (Supplementary Fig. S5g-i).

Whilst derived from just two of the many neuronal subtypes in the fly nervous system, these data indicate that GOF BK channels do not grossly disrupt neural arborization and synaptic growth in *Drosophila* – a conclusion consistent with our finding that the number and size of presynaptic boutons at the neuromuscular junction are unchanged in *slo*^E366G/+^ larvae^26^. We therefore focused our investigations on the final stages of synaptic maturation, during which GOF BK channels exert their pathological effect on movement (Fig. 3b) ^37^. In late-stage *slo*^E366G/+^ pupae, immuno-fluorescent imaging revealed an ∼ 35% reduction in brain-wide synaptic localisation of Bruchpilot (BRP), a key presynaptic scaffold protein that tethers synaptic vesicles in proximity to calcium channels within the active zone ^42^ (Fig. 6a-c). Intriguingly, the reduction in BRP localisation that occurred during the late pupal stage partially persisted into adulthood (Fig. 6b, c). In contrast, during the mid-pupal stage, when expression of GOF BK channels minimally effects movement (Fig. 3b), synaptic BRP localisation was similar between *slo*^E366G/+^ and controls (Fig. 6b, c). These data suggest that BK channel GOF leads to changes in presynaptic composition during neurodevelopment that partially persists into adulthood.

**Fig. 6.**
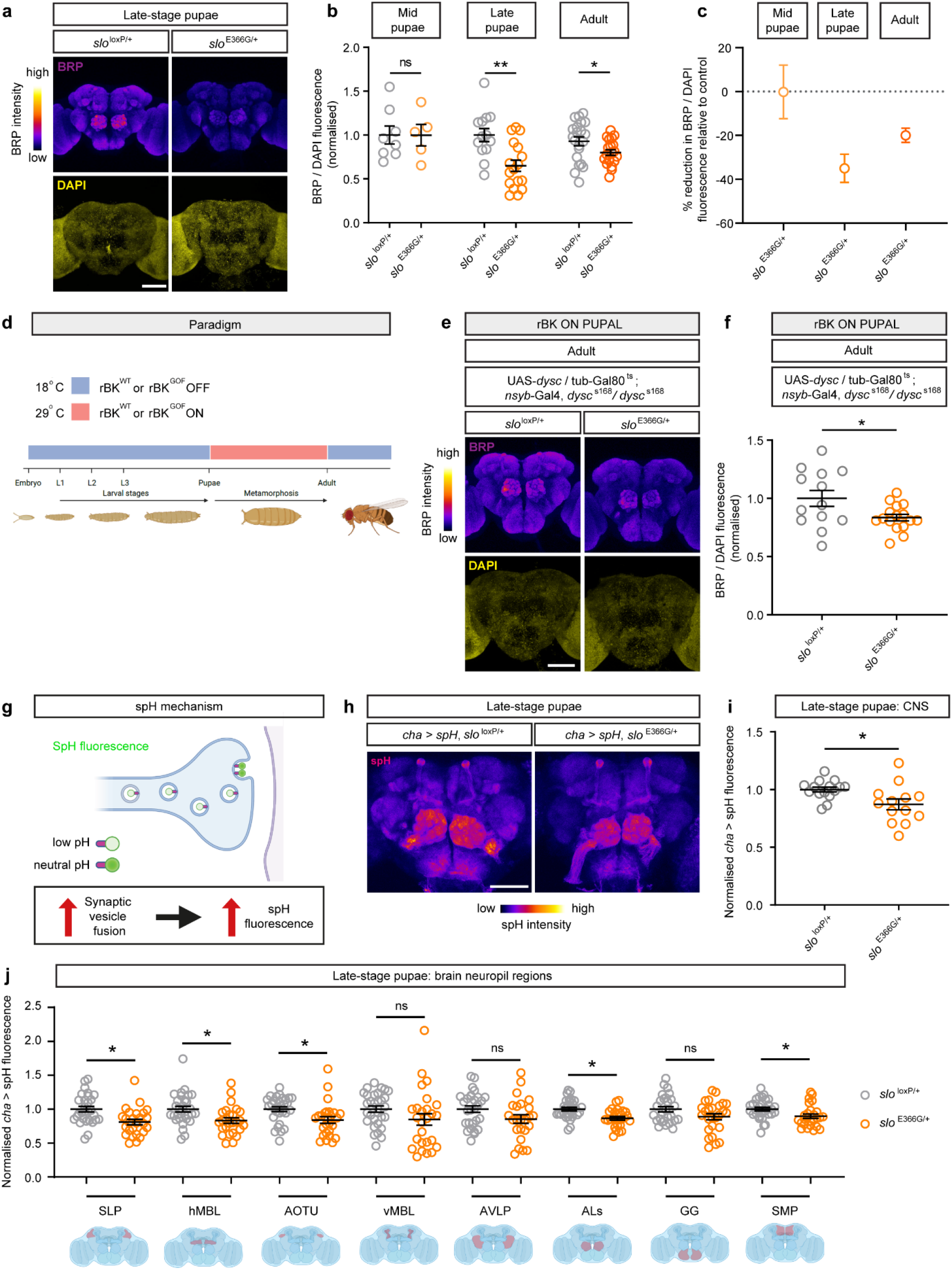
BK channel GOF reduces Bruchpilot expression and excitatory neurotransmission during late neurodevelopment. **(a)** Representative images of control (*slo*^loxP/+^) and BK channel GOF (*slo*^E366G/+^) late-stage pupal ex vivo brains stained with anti-Bruchpilot (BRP) and counterstained with DAPI. **(b)** Quantification of whole-brain BRP fluorescence normalised to DAPI counterstain, at the P8-9 mid-pupal stage (n = 8, 5 brains), P14 late-pupal stage (n = 13, 17 brains), and in 5-7 d post-eclosure adults (n = 21, 20 brains) in *slo*^loxP/+^ and slo^E366G/+^ flies. Data are presented normalised to the corresponding *slo*^loxP/+^ mean. **(c)** Data from (b) represented as % change from the relevant *slo*^loxP/+^ control mean. **(d)** Schematic illustrating paradigm to induce neuronal rBK^WT^ or rBK^GOF^ expression during the pupal stage. Brains were removed and visualised ex vivo at the p14 late-pupal stage. **(e)** Representative images of control and BK channel GOF late-stage pupal ex vivo brains immuno-stained with anti-BRP and counterstained with DAPI, in the paradigm illustrated in (d). **(f)** Quantification of whole-brain BRP fluorescence, normalised to DAPI counterstain in *slo*^loxP/+^ (n = 13 brains) and slo^E366G/+^ (n = 16 brains) late-stage ex vivo pupal brains in the paradigm illustrated in (d). Data are presented normalised to the control mean. **(g)** Schematic illustrating mode of action of synaptopHluorin (spH). Enhanced fluorescence is indicative of increased synaptic vesicle release. **(h)** Representative images of spH expressed in excitatory cholinergic neurons using the *cha-* Gal4 driver in control (*cha > spH*, *slo*^loxP/+^) and BK channel GOF (*cha*>*spH*, *slo*^E366G/+^) late-stage pupal ex vivo brains. **(i)** Quantification of whole-brain spH fluorescence in *slo*^loxP/+^ control (n = 14 brains) and *slo*^E366G/^ BK channel GOF (n = 13 brains) the central region of late-stage pupal ex vivo brains. Data are presented normalised to the control mean. **(j)** Quantification of spH fluorescence across neuropil domains in control (*slo*^loxP/+^) and BK channel GOF (*slo*^E366G/+^) ex vivo late-stage pupal brains. Data are presented normalised to *slo*^loxP/+^ means within each neuropil domain. The location of each domain within the late pupal brain is shown schematically below each dataset. Each datapoint represents data from a given neuropil domain with an individual hemisphere. Abbreviations: SLP – super lateral protocerebrum, hMBL – horizonal mushroom body lobes; AOTU – anterior optic tubercle; vMBL – vertical mushroom body lobes; AVLP – anterior ventrolateral protocerebrum; ALs – antennal lobes; GG – gnathal ganglia; SMP – superior medial protocerebrum. n = 26-30 per neuropil domain. Error bars: SEM. *p < 0.05, **p < 0.005, ***p < 0.0005, ns – p > 0.05, t-test with Welch’s correction or Mann-Whitney U-test for parametric and non-parametric datasets respectively.

We sought to probe the mechanisms underlying reduced synaptic BRP localisation in *slo*^E366G/+^ late pupae and adults. Induction of neuronal rBK^GOF^ expression solely during the pupal stage significantly reduced synaptic BRP localisation in resulting adult brains (Fig. 6d-f), indicating that GOF BK channels act during neurodevelopment to influence adult synaptic BRP localisation. Surprisingly, western blotting revealed that BRP protein levels showed a small (∼ 15%), marginally significant increase in *slo*^E366G/+^ compared to control late-stage pupal brains (Supplementary Fig. S6a, b). Additionally, the localisation of Discs Large (DLG) – a post-synaptic scaffold protein that, like BRP, is pan-neuronally expressed in the *Drosophila* brain ^43^ – was unaltered in late-stage *slo*^E366G/+^ pupal brains compared to controls (Supplementary Fig. S6c, d). These data suggest that the reduction in synaptic BRP localisation in *slo*^E366G/+^ late pupal brains is not driven by reduced BRP protein levels or synapse/neuronal loss, but may be indicative of a decrease in active zone number or altered distribution of BRP within presynaptic domains ^44^ (see Discussion).

### BK channel GOF suppresses excitatory neurotransmission during development

Developing nervous systems, including that of *Drosophila*, are characterised by stimulus-independent electrical activity that facilitates synaptic connectivity via Ca^2+^-dependent signalling cascades ^24,45–47^. Since BRP promotes neurotransmitter release at the *Drosophila* larval NMJ ^42^, reductions in presynaptic BRP levels in *slo*^E366G/+^ late-stage pupae might inhibit neurotransmitter release during this period, as could enhanced activity of BK channels localised to presynaptic domains ^8^. Hence, reducing neurotransmission during neurodevelopment represents a plausible mechanism by which BK channel GOF could disrupt adult-stage motor control.

We therefore tested whether BK channel GOF reduced neurotransmission during the late pupal stage of *Drosophila*. To do so, we used synaptopHluorin (spH), a pH-sensitive GFP variant localised to the lumen of synaptic vesicles ^48^. Fusion of acidified vesicles with the presynaptic membrane results in local pH neutralisation and an increase in spH fluorescence, providing an optical readout of neurotransmitter release (Fig. 6g) ^48^. Since excitatory transmission promotes neuronal Ca^2+^ influxes during development, and acetylcholine is the predominant excitatory neurotransmitter in insects, we expressed spH in cholinergic neurons and imaged spH fluorescence in control and *slo*^E366G/+^ late-stage pupal brains, using an ex vivo preparation to visualise neurotransmission occurring independently of sensory input. As predicted, we observed a significant reduction in spH fluorescence across cholinergic synapses in *slo*^E366G/+^ late-stage pupal brains compared to controls (Fig. 6h, i). Comparisons of cholinergic spH fluorescence in specific neuropil domains revealed that this reduction encompassed several brain regions that regulate movement and which are activated prior to or during locomotion, including the anterior optic tubercle, superior medial protocerebrum, and the mushroom bodies ^49–52^ (Fig. 6j). Thus, BK channel GOF widely inhibits excitatory neurotransmission during a critical late-stage period of *Drosophila* neurodevelopment.

### Reduced neural activity during development disrupts limb control in *Drosophila*

Is reducing neurotransmission in the developing fly brain sufficient to cause motor dysfunction in adult flies? To examine this question, we expressed *dOrk*ΔC2 – a constitutively active open rectifying channel that reduces neuronal excitability by hyperpolarising the resting membrane potential ^53^ – solely in pupal post-mitotic neurons and measured the impact on overall movement and limb kinematics in resulting adult flies (Fig. 7a, b). We found that a large percentage of flies subjected to pupal pan-neuronal silencing failed to eclose from the pupal case (data not shown), suggestive of extreme motor dysfunction. Using the DAM system, we indeed found that flies that survived this developmental manipulation exhibited a significant reduction in movement compared to flies of the same genotype kept at restrictive, non-silenced conditions (Fig. 7b). Experiments in control flies demonstrated that the temperature shift required to limit neuronal dOrkΔC2 expression to the pupal stage did not account for this effect (Fig. 7b). We next used the FLLIT system to test how limb kinematics were altered by pupal-stage neuronal silencing (Fig. 7c-h). Strikingly, three of four aspects of limb movements impaired in *slo*^E366G/+^ flies (the % of time each limb moved, displacement, and non-linearity of limb movement; but not limb velocity) were also perturbed by neuronal silencing during development (Fig. 7e-h).

**Fig. 7.**
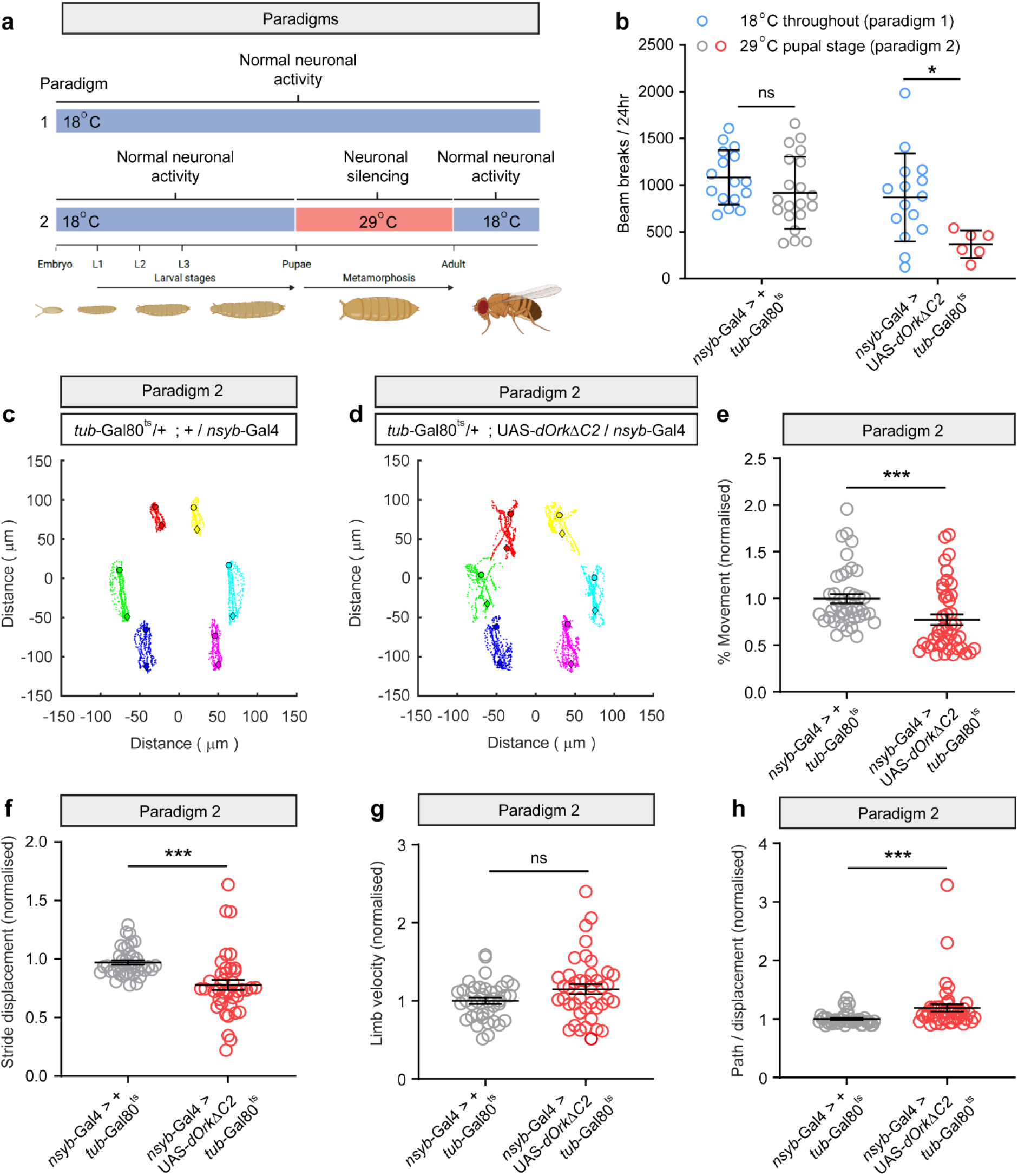
Suppressing neuronal excitability during late development disrupts locomotor activity and limb kinematics. **(a)** Schematics illustrating the temporal pan-neuronal expression of OrkΔC2, which hyperpolarises and silences neurons. OrkΔC2 expression during the pupal stage was terminated shortly before eclosure (paradigm 2). Experiments on adult male flies took place 5-7 days after eclosure. **(b)** Total number of beam breaks in a 24 h period recorded using the DAM system in the paradigms illustrated in (a). *tub-*Gal80^ts^, *nsyb-*Gal4 */ +*: paradigm 1, n = 16; paradigm 2, n = 20. *tub-*Gal80^ts^, *nsyb-*Gal4 > UAS-*OrkΔC2*: paradigm 1, n = 15; paradigm 2, n = 6. **(c-d)** Representative FLLIT-derived traces showing movement of each limb over 3-6 strides, relative to the centre of the body, for the control (*tub-*Gal80^ts^, *nsyb-*Gal4 */ +*) (c) and experimental (*tub-*Gal80^ts^, *nsyb-*Gal4 > UAS-*OrkΔC2*) (d) flies subject to experimental paradigm 2. **(e-h)** FLLIT-derived gait parameters comparing control to experimental flies. Data points represent mean values from a single limb across 3-6 strides. Data are presented normalised to control means. n = 42 limbs across 7 flies for both genotypes. Error bars: SEM. *p<0.05, **p < 0.005, *** p< 0.0005, ns – p> 0.05, Mann-Whitney U-test (e-h) or chi-squared test (i).

Furthermore, we noted an increase in the variability of limb displacement and movement linearity in flies subjected to pupal neuronal silencing (Supplementary Fig. S7a-c), as was observed in in *slo*^E366G/+^ flies and in flies with neuronal rBK^GOF^ expression induced during the late pupal stage (Supplementary Fig. S4). Thus, reduced neuronal excitability during neurodevelopment, as is observed in *slo*^E366G/+^ pupae, is sufficient to impair movement in resulting adult flies by altering specific aspects of limb kinematics and movement stereotypy.

### Elevating neural activity during development rescues motor defects caused by BK channel GOF

Finally, we sought to demonstrate a causal link between reduced neurotransmission during neurodevelopment and disrupted movement and limb kinematics in *slo*^E366G/+^ flies. To do so, we tested whether increasing the excitability of pupal neurons could rescue movement defects in *slo*^E366G/+^ adults. We deployed TrpA1 – a thermo-sensitive cation channel mildly activated at 27°C and strongly at 29°C ^54^ – to elevate the activity of defined pupal neurons in *slo*^E366G/+^ and control backgrounds. We initially induced strong elevations in the activity of all cholinergic neurons in *slo*^E366G/+^ and control pupae, but this manipulation was largely lethal in the *slo*^E366G/+^ background, with very few flies eclosing from the pupal case. Since this suggested that BK GOF flies were particularly sensitive to the levels of neural activity during development, we induced a more subtle manipulation of neural activity. Recent work has identified a small subpopulation of neurons defined by expression of the Trp-γ cation channel which promote stimulus-independent neural activity during development ^55^. Trp-γ-neuron axons ramify broadly throughout the developing brain and this cell-type has been hypothesised to act as a relay network that transmits excitatory signals from a putative central pattern generator to the wider nervous system during the pupal stage ^55^ (Supplementary Fig. S8a). We used a previously-defined reporter of Trp-γ expression ^56,57^ that labels neurons in the gnathal ganglia, optic lobes, and superior protocerebral domains of the pupal brain (Supplementary Fig. S8b, c). We used this reporter to drive TrpA1 expression in Trp-γ-neurons; induced mild or strong activation of TrpA1 in control or *slo*^E366G/+^ pupae via changes in ambient temperature (Fig. 8a); and quantified overall movement in resulting adults using the DAM system (Fig. 8b). In *slo*^loxP/+^ controls, we observed a detrimental effect of high temperature during the pupal stage on adult movement, with no specific impact of activation of Trp-γ-neurons being apparent (Fig. 8b). In contrast, in the *slo*^E366G/+^ background, both mild and strong elevation of Trp-γ-neuron activity significantly enhanced overall movement compared to *slo*^E366G/+^ flies expressing TrpA1 channels that were inactive during neurodevelopment, with strong activation of Trp-γ-neurons during development inducing the most robust rescue (Fig. 8b).

**Fig. 8.**
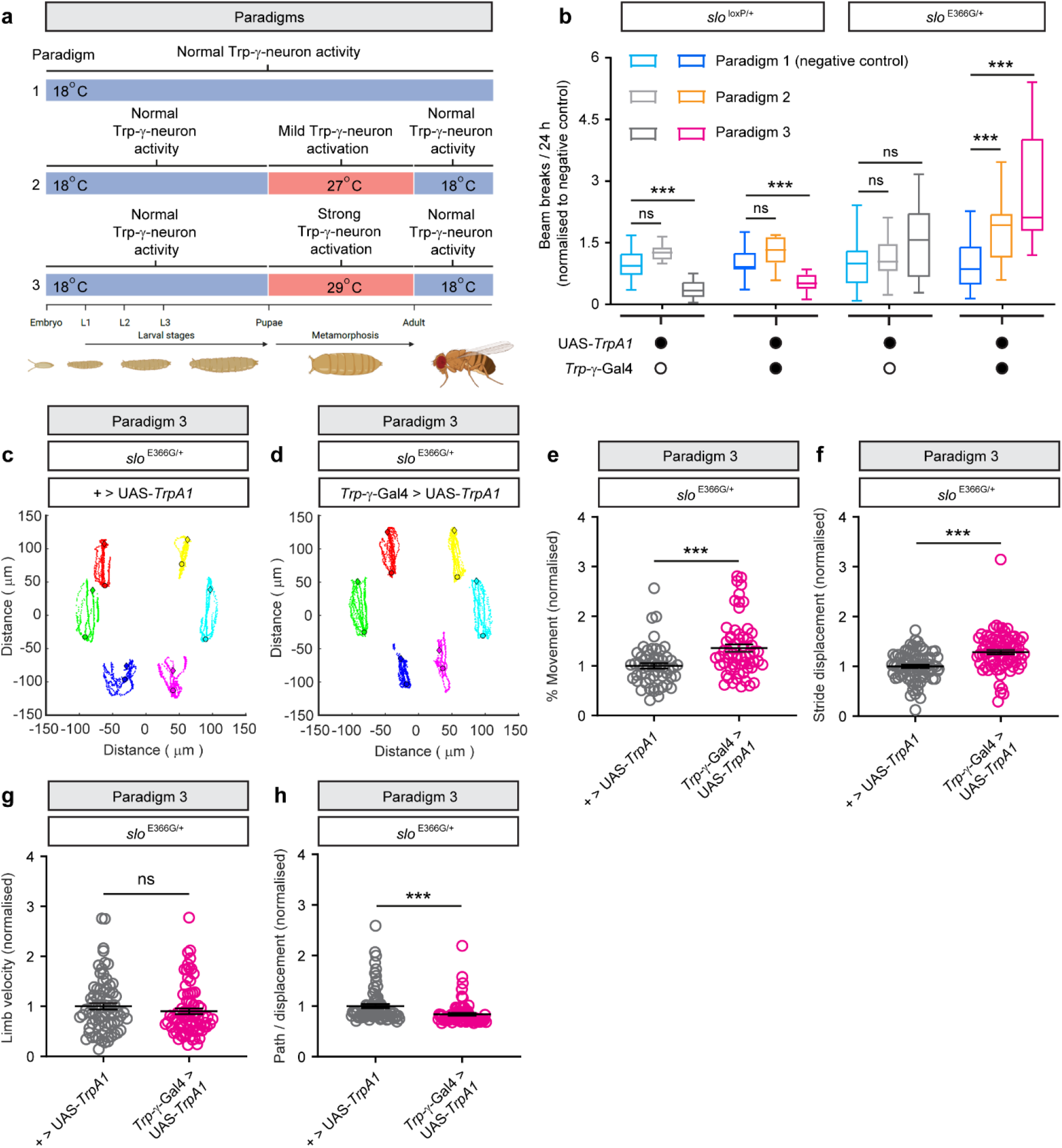
Enhancing neuronal excitability during development partially rescues locomotor effects caused by BK channel GOF. **(a)** Schematics illustrating the activation of Trp-γ neurons using the heat-sensitive cation channel TrpA1. Mild (paradigm 2) or strong (paradigm 3) Trp-γ neuron activation during the pupal stage was terminated shortly before eclosure. Experiments were performed at 5-7 d post-eclosure. **(b)** Total beam breaks over a 24 h period, recorded using the DAM system, in control (*slo*^loxP/+^) or BK channel GOF (*slo*^E366G/+^) flies subject to the paradigms illustrated in (a). UAS- *TrpA1 / +*, *slo*^loxP/+^: n = (left to right) 24, 6, and 20 flies. *Trp-γ* > UAS-*TrpA1, slo*^loxP/+^: n = (left to right) 27, 11, 10. UAS-*TrpA1 / +*, *slo*^E366G/+^: n = (left to right) 47, 28, 6. *Trp-γ* > UAS-*TrpA1, slo*^E366G/+^: n = (left to right) 49, 19, 9. Data were normalised to the mean of the relevant genotype in paradigm 1. Data are presented as Tukey box plots. Box denotes the median, 25% and 75% percentiles. **(c-d)** Representative FLLIT-derived traces showing movement of each limb over 3-6 strides, relative to the centre of the body, for flies expressing GOF BK channels (slo^E366G/+^) without (UAS-*TrpA1 / +*) (c) and with (*Trp-γ* > UAS-*TrpA1*) (d) strong excitation of Trp-γ neurons via TrpA1 activation via experimental paradigm 3. (**e-h**) FLLIT-derived gait parameters in the above backgrounds subject to experimental paradigm 3. Data points represent mean values from a single limb across 3-6 strides. Data are presented normalised to control means. n = 78 limbs across 13 flies for both genotypes. Error bars: SEM. *p<0.05, *** p< 0.0005, ns – p> 0.05, two-way ANOVA (b) or Mann-Whitney U-test (e-h).

We next tested how strong Trp-γ-neuron activation during the pupal stage affected limb kinematics and dyskinesia-like limb movements in *slo*^E366G/+^ flies. Using the FLLIT system once more, we found that three out of the four kinematic parameters perturbed in *slo*^E366G/+^ flies were partially rescued, with pupal Trp-γ-neuron activation inducing an increase in the % time each limb was moving, an increase in stride displacement, and an increase in the linearity of limb movement (Fig. 8c-f, h; compare with Fig. 4d-g). Notably, these same kinematic parameters were disrupted by neuronal silencing during the pupal stage (Fig. 7e, f, h), strongly indicating that they are influenced by neural activity during development. In contrast, the limb velocity of *slo*^E366G/+^ flies – a kinematic parameter unaffected by neural silencing during the pupal stage (Fig. 7g) – was unaltered by pupal Trp-γ-neuron activation (Fig. 8g). Nor did we observe a substantial reduction in the variability of limb displacement, the non-linearity of limb movement, or the number or duration of dyskinesia-like leg twitches, in *slo*^E366G/+^ flies following activation of pupal Trp-γ-neuron (Supplementary Fig. S8d-h).

Collectively, the above data uncover specific components of the multi-faceted motor phenotype of *slo*^E366G/+^ flies that are sensitive or insensitive to neuronal activity during the development of the adult *Drosophila* nervous system.

## Discussion

Our data define a critical developmental window in which GOF BK channels act to perturb movement in *Drosophila*; uncover alterations in synaptic protein expression and neurotransmission during this critical period; and delineate causative relationships between specific aspects of adult *Drosophila* limb movements and the levels of neural activity occurring during neurodevelopment.

Our core results are derived from a method of promoting or inhibiting robust BK channel expression involving spatio-temporal manipulation of the SLO-binding protein DYSC^27^. A caveat of this method is that behavioural experiments are performed in a *dysc* mutant background. Since *dysc* mutants exhibit defects in circadian patterns of locomotion ^27^ and fly movement was measured by the DAM system over a 24 h period, alterations in circadian activity could potentially yield unforeseen confounds. We control for this using two video-based methods that measure movement capacity and limb kinematics independently of circadian fluctuations in movement: the DART and FLLIT systems ^36,39^. Importantly, data derived from these hi-resolution methods support a pathological impact of GOF BK channels during development (Fig. 2 and Fig. 4).

While we were able to use GAL80^ts^ to increase expression of BK channels in pupal neurons (Supplementary Fig. S1), the degradation rate of DYSC and SLO proteins are also unknown. Thus, a second caveat of our approach is that BK channels induced during the pupal stage may be partially retained into adulthood. Hence, we cannot fully rule out a pathogenic effect of GOF BK channels in adult neurons. Nonetheless, observations from *slo*^E366G/+^ flies (which are wild-type except for a single heterozygous mis-sense mutation in *slo*) support a neurodevelopmental impact of GOF BK channels. Firstly, abnormal limb kinematics in *slo*^E366G/+^ flies are similar to those caused by limiting neuronal expression of GOF BK channels to the pupal stage via temporal control of DYSC (Fig. 4 and 5). Secondly, we observed a reduction in presynaptic BRP in *slo*^E366G/+^ neurons that emerged during, but not before, the critical period identified by behavioural analyses of flies subject to dynamic alterations in DYSC and SLO expression (Fig. 6). Thirdly, we identify reductions in excitatory neurotransmission in *slo*^E366G/+^ pupal neurons and found that artificially reducing neural activity of pupal neurons in a wild-type background partially phenocopied the effect of BK channel GOF on locomotor activity and limb kinematics (Fig. 7). Fourthly, enhancing neural activity solely during the pupal stage partially rescued motor defects in *slo*^E366G/+^ adult flies (Fig. 8). Since this intervention does not rely on protein translation and degradation but instead involves rapid activation/inactivation of TrpA1 cation channels ^54^, direct effects on neuronal excitability in the adult nervous system can be excluded.

These complementary datasets thus support our core conclusion: that GOF BK channels impair limb control in *Drosophila* by perturbing neuronal development, an effect that occurs in part through inhibition of neural activity in the developing brain.

### BK channels and neurodevelopment

A neurodevelopmental impact of GOF BK channels is consistent with studies of humans harbouring BK channel GOF mutations. Human brain development extends into the teenage years ^58^, and paroxysmal dyskinesia human hSlo1 D434G carriers initiates mainly before the age of 6 (in some cases less than 6 months after birth) ^12^. Hence, key pathogenic processes that cause loss of limb control in hSlo1 D434G carriers occur during neurodevelopment. Furthermore, while hSlo1 D434G carriers do not present with overt neurodevelopmental phenotypes, hSlo1 mutations such as N999S that enhance BK channel activity to a greater degree than D434G are associated with neurodevelopmental morbidities such as developmental delay, intellectual disability, and autism spectrum disorder ^21,59,60^.

In concert with these studies, our findings suggest a model in which subtle alterations in neurodevelopment caused by mild BK channel GOF are sufficient to disrupt limb control, while greater enhancement of BK channel activity further disrupts critical neurodevelopmental processes to an extent that yields neurodevelopmental pathologies. Interestingly, a comprehensive study by Dong et al., recently identified locomotor defects and dyskinesia-like movements in knock-in mice carrying the equivalent *KCNMA1* D434G allele ^19^. Notably, locomotor defects in *KCNMA1* D434G mice were partially rescued by intraperitoneal injection of Paxilline, a BK channel blocker. While these results imply an acute role for BK GOF in causing locomotor defects, our results suggest an alternative interpretation: that modulating action potential frequency or neurotransmission via BK channel inhibition may counteract the effect of prior neurodevelopmental perturbations caused by BK GOF. Whether Paxilline rescues dyskinesia-like phenotypes in *KCNMA1* D434G mice also remains unclear. Notably, Dong et al. identified alterations in the somatic and dendritic morphology of cerebellar Purkinje neurons in *KCNMA1* D434G mice ^19^, supporting a link between BK GOF and altered neurodevelopment in mammals. An important extension of our study will therefore be to develop genetic methods to induce GOF BK expression in the developing or developed murine nervous system in BK GOF mouse models ^19,20^ and define the resulting impact on locomotor and dyskinesia-like phenotypes.

### A complex interplay between BK channels, neurotransmission, and synaptic maturation

Numerous studies have demonstrated a critical role for neural activity during development in regulating synaptic connectivity ^24,61,62^, providing a plausible link between BK channel GOF, excitatory neurotransmission, and an altered developmental state of pre-motor circuits that disrupts limb control. In support of this notion, stimulus-independent neural activity has been detected in the vertebrate developing spinal cord, and suppression of this activity results in a variety of defects in pre-motor circuit development and movement, including axonal mis-wiring ^63^, delayed onset of co-ordinated movement ^64^, and aberrant patterns of muscle activation ^65^. Disrupting neural activity in *Drosophila* embryos similarly causes long-lasting defects in the crawling behaviour of resulting larvae ^66,67^.

However, further work is required to examine how excitatory neurotransmission, synaptic protein localisation, and synaptic connectivity, are perturbed by BK GOF. Super-resolution imaging has revealed that loss of SLO BK channels enlarges BRP-labelled active zones at the larval *Drosophila* NMJ ^68^, and that altered compaction of active zones can change the immuno-fluorescence of their protein constituents when visualised through confocal microscopy ^44^. Such a phenomenon could explain why BRP immuno-fluorescence is reduced in pupal *slo*^E366G/+^ neurons despite BRP protein levels not decreasing (Fig. 6 and Supplementary Fig. S6). Alternatively, changes in the trafficking of BRP or a decrease in the number of presynaptic active zones could drive reduced BRP immuno-fluorescence in pupal *slo*^E366G/+^ neurons. One future avenue of research will therefore be to define in more detail how the structure of the active zone cytomatrix is modified by BK GOF through techniques such as super-resolution imaging or electron microscopy.

A second line of investigation will be to disentangle the relationship between GOF BK channels, neural activity, and active zone protein localisation. Presynaptic BK channels can negatively tune neurotransmission by inactivating presynaptic voltage-gated calcium channels ^8^, and a reduction in BRP levels would also be expected to reduce synaptic release^42^. However, suppressing neural activity during development through Trp-γ-neuron silencing predominantly reduces the number of BRP-labelled active zones in neurons of the fly visual system ^55^, illustrating a bi-directional relationship between neural activity and synaptic active zone structure during development. Thus, whether GOF BK channels influence BRP localisation by reducing neurotransmission, or instead regulate BRP through activity-independent mechanisms to then reduce excitatory neurotransmission, remains unclear.

It is also important to note that not all aspects of altered movement and limb control in *slo*^E366G/+^ flies were modulated by enhancing neural excitability during development. As well as neuronal plasma membranes, BK channels are localised to the nuclear envelope, mitochondria, and lysosomes, where they control intra-organellar Ca^2+^ levels ^69–71^. Hence, BK GOF may cause specific impairments in limb control, such as the dyskinesia-like movements observed in *slo*^E366G/+^ flies, by disrupting processes such as gene expression, metabolism, or protein degradation during neurodevelopment. The generation of invertebrate and vertebrate animal models harbouring BK channel GOF mutations ^19,20,26^ now provides a means to test this hypothesis.

### Critical windows in involuntary movement disorders

Recent work has revealed that neurological disorders lacking overt neurodevelopmental phenotypes may nonetheless have important developmental components. For example, human foetuses carrying pathogenic HTT alleles linked to Huntington’s Disease (HD), a prototypical age-related disorder associated with involuntary choreiform movements, exhibit anomalies in cortical development ^72^. Furthermore, mouse models of HD exhibit reduced excitatory cortical activity specifically in the first post-natal week, and pharmacologically elevating activity during this early period rescues sensorimotor defects in HD mice ^73^. Elegant work has similarly identified an early critical therapeutic window in the context of isolated dystonia caused by a LOF mutation in the *TOR1A* AAA+ ATPase, showing that restoration of *TOR1A* expression in juvenile but not adult neurons rescued dystonia-like phenotypes in *TOR1A* LOF mice ^74^. Studies of *Drosophila* have further supported the concept that early perturbations in the excitability of developing circuits can lead to long-lasting changes in behaviour and nervous system robustness ^75,76^. Our work further emphasises the potential contribution of developmental alterations to involuntary movement disorders lacking overt neurodevelopmental phenotypes, and reveals the existence of a critical upper limit on developmental BK channel activity that, when exceeded, disrupts limb control.

## ACKNOWLEDGEMENTS

We thank Prof. Kyunghee Koh (Thomas Jefferson University), Prof. Gero Miesenböck (University of Oxford), and Prof. Nicolas Tapon (Francis Crick Institute) for kindly sharing *Drosophila* stocks. We thank Dr. Amanda Pocratsky (UCL Queen Square Institute of Neurology, UK) for comments on the manuscript. This study was funded by a DMRF Post-Doctoral Fellowship to Dr. Simon Lowe; a Wellcome PhD studentship award to Dr. Patrick Kratschmer (109003/Z/15/Z); a National Medical Research Council Open Fund Individual Research Grant (OFIRG19nov-0063) to Dr. Sherry Aw; and a MRC Senior Non-Clinical Fellowship (MR/V03118X/1), BBSRC Project Grant (BB/X00094X/1), and Wellcome Strategic Award (WT104033AIA) to Prof. James Jepson. Schematics shown in Figures 1a, 1b, 1d, 2b, 3a, 4a, 5a, 5h, 6d, 6g, 6j, 7a, 8a; and Supplementary Figures S1b, S1e, S2a, S2c, S3a, S3d, S4c, S4f, S7a, S8d; were created with Biorender.com.

## AUTHOR CONTRIBUTIONS

S.A.L – Conceptualization, Methodology, Investigation, Writing - Original Draft, Writing – Review & Editing, Visualization, Funding acquisition; A.D.W – Investigation, Visualization, Writing – Review & Editing; G.A – Software, Formal analysis, Writing – Review & Editing; A.B – Investigation, Software, Resources, Writing – Review & Editing; T.G – Investigation, Writing – Review & Editing; N.S-B – Investigation; A.S – Investigation, Writing – Review & Editing; M.B – Investigation; A.G – Investigation; P.K – Resources; S.S.A – Resources, Software, Writing – Review & Editing Supervision, Funding acquisition; J.E.C.J – Conceptualization, Investigation, Visualization, Writing - Original Draft, Writing – Review & Editing, Supervision, Project administration, Funding acquisition.

## Materials and Methods

### *Drosophila* husbandry

Flies were maintained on standard fly food under 12 h: 12 h light-dark cycles (12L: 12D). Unless otherwise indicated flies were maintained at a constant 25°C. For temperature manipulations they were maintained at 18°C, 27°C or 29-30°C at the time stages indicated. To control for genetic background all lines used were backcrossed into the isogenic w^1118^ (iso31) background for at least 5 generations. Genotypes used were:

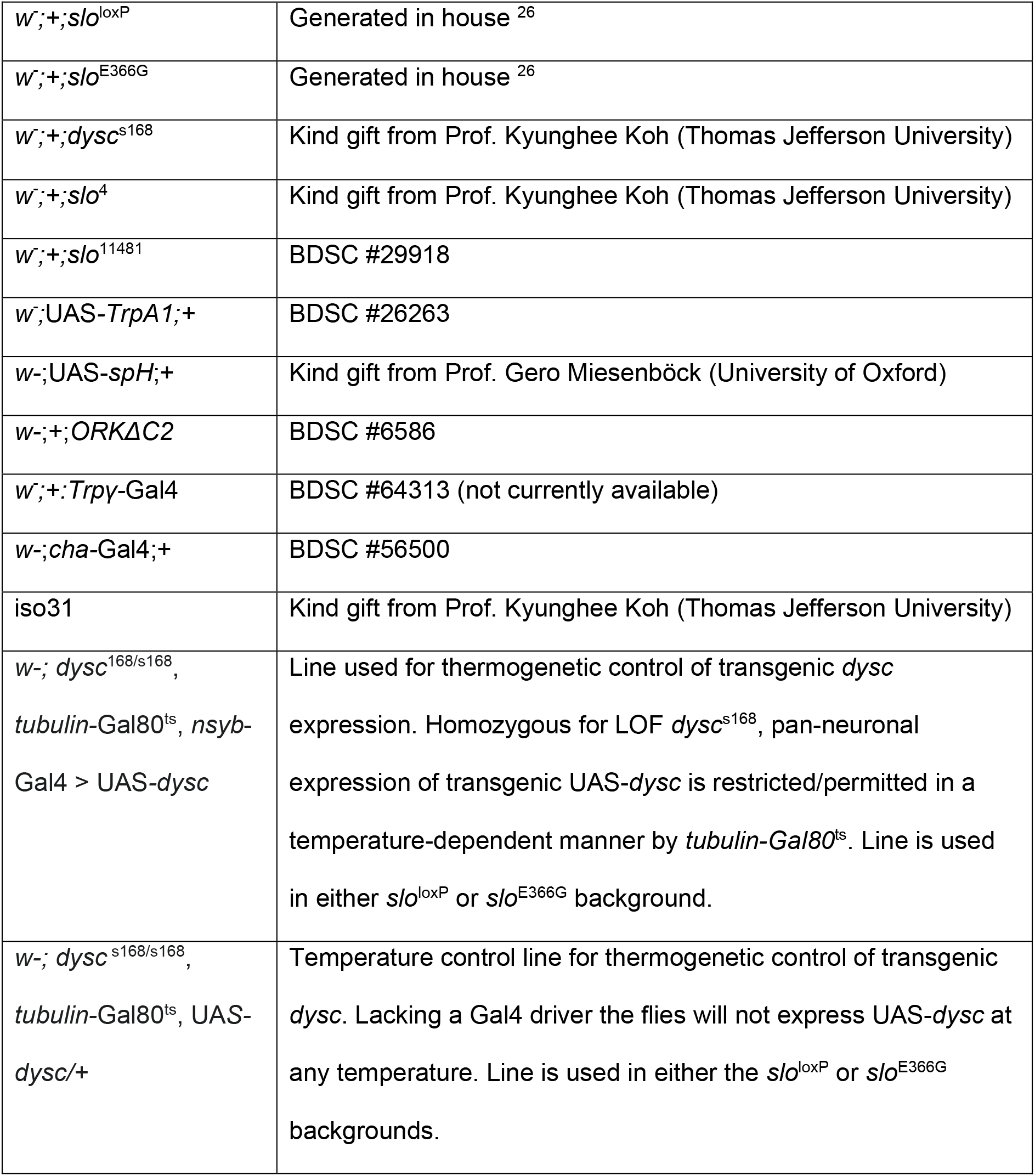
w- indicates either w^1^ or w^1118^.

### Antibodies and western blotting

Lysates were prepared from 10 homogenised adult or pharate brains in lysis buffer with a phosphatase–protease inhibitor cocktail (Roche) on ice and suspended in LDS sample buffer and reducing agent (Invitrogen). Samples were resolved on a 4-12% polyacrylamide gel (Bolt Bis-Tris Plus, Invitrogen) and transferred to a PVDF membrane. Membranes were blocked for 1 hr at RT in 3% bovine serum albumin and 5% low-fat milk in Tris-buffered saline and 0.1% Tween-20 (TBS-T). Primary antibodies for SLO (1:500 PA1-923, Invitrogen) or BRP (1:100 nc82, DSHB) were incubated overnight at 4°C. Membranes were washed with TBS-T and incubated with goat anti-rabbit/mouse HRP-conjugated secondary antibody (Invitrogen). Antibody-antigen complexes were visualised by SuperSignal West Pico Plus chemiluminescent substrate (Thermo Scientific) with an iBright 750 Imaging System (Invitrogen). For loading controls, total protein was quantified per lane with a Novex reversible membrane protein stain (Invitrogen). SLO protein expression was normalised to a non-specific band present in the *slo* null sample. Densities of proteins were quantified on ImageLab 6.1 software.

### Immuno-histochemistry

Brains were dissected in phosphate-buffered saline (PBS) (Sigma Aldrich) at the time point indicated (3-7 day old adult, pharate adult [p14] or mid-pupal stage [p8-9]) and immuno-stained as described previously ^77^. Briefly, brains were fixed by 20 min room temperature incubation in 4% paraformaldehyde (MP biomedicals), and blocked for one hour in 5% normal goat serum in PBS+0.1% Triton-X (Sigma-Aldrich) (PBT). Brains were incubated overnight in primary antibody at 4°C, washed three times in PBT, and incubated overnight in secondary antibody at 4°C. Primary antibodies use were: mouse anti-BRP/nc82 at 1:500, mouse anti-Unc18/Rop 4F8 at 1:500, mouse anti-discs large at 1:1000, mouse anti-PDF at 1:100, mouse anti-FASII at 1:200 (all from Developmental Studies Hybridoma Bank, University of Iowa, Iowa City, IA, USA; AB Registry ID: AB_528484). Secondary antibodies were goat anti-mouse AlexaFluor 555, 568, and 647 at 1:1000, and goat anti-rabbit 555 at 1:1000 (Life Technologies). Counterstain was DAPI at 1:1000 (ThermoFisher Scientific). Brains were mounted and imaged in SlowFade Gold anti-fade mountant (Thermofisher). Images were taken with a Zeiss LSM 710 confocal microscope with an EC ‘Plan-Neofluar’ 20x/0.50 M27 air objective, taking z-stacks through the entire brain with step sizes of 1-5 μm. Images were analysed using ImageJ: z-stacks were 3-D projected using a maximum intensity projection, ROIs were drawn around the central brain (excluding optic lobes), mean fluorescence values were taken with background subtracted, and values were normalised against mean fluorescence of the counterstain. Images were compared to controls imaged on the same day, and normalised to the mean values of controls.

### Live imaging

Transgenic synapto-pHluorin (UAS-*spH*) was expressed in cholinergic neurons using the *cha*-Gal4 driver. Pharate adult (p14) brains were dissected in external solution (in mM: 101 NaCl, 1 CaCl_2_, 4 MgCl_2_, 3 KCl, 5 glucose, 1.25 NaH_2_PO_4_, 20.7 NaHCO_3_, pH 7.2). Images were taken immediately (within 5 min) as above. ROIs were drawn around the whole central brain as above, or around neuropil structures identifiable within the image, backgrounds subtracted, and mean fluorescence values taken. Values were normalised to the mean values of controls taken during the same session.

### Adult behavioural analyses

#### DAM

*Drosophila* Activity Monitor (DAM) analysis was performed as described previously ^26^. Briefly, three- to five-day old male flies were collected and loaded into glass behavioural tubes (Trikinetics inc., MA, USA) containing 4% sucrose and 2% agar and left for two full days to acclimatise at the relevant ambient temperature. On the third day the total number of beam breaks over 24 h was measured using *Drosophila* Activity Monitor (DAM, Trikinetics inc., MA, USA). Experiments were conducted in in 12L:12D at the indicated temperature.

#### DART

Five- to seven-day old male flies were loaded into behavioural tubes and maintained in conditions similarly to above, but with flies maintained for 24 h at the relevant temperature before mechanical stimuli were applied. Locomotor activity was recorded using the *Drosophila* ARousal Tracking (DART) system (BFKlabs) ^36^ as previously described ^26^. After 24 h acclimatisation, a single stimulus in the form of a mechanical vibration (a train of five 200 ms pulses separated by 800 ms intervals, set at the DART system’s maximum intensity) was delivered at Zeitgeber Time 3 (ZT3), causing a robust locomotor response. Videos were taken using a USB webcam (Logitech) and speed (mm/s) was analysed by the DART system from absolute position, and binned in one min intervals. Flies were excluded from analysis if they exceeded an average of 1 mm / s speed in the five one-minute bins preceding the stimulus. The mean speed (mm/s) in the one-minute bin following the stimulus was analysed.

#### FLLIT

For FLLIT (Feature Learning-based LImb segmentation and Tracking) measurements, five- to seven-day old de-winged male flies were loaded into custom-made 2×2 cm arenas without anaesthesia. 1000 fps videos consisting of flies walking in a straight line and taking at least three clear strides were taken using a Photron FASTCAM Mini UX50 High Speed Camera with a Sigma 105 mm Macro lens. Gait analysis was performed using the FLLIT system ^3^.

Where FLLIT generates data for parameters per stride, the first and last stride were excluded and a mean of the remaining values taken. Values for each leg were pooled, giving six values per fly.

Table of parameters and definitions

| <u>Parameter</u> | <u>Calculation</u> |
| --- | --- |
| Movement % | % of the video for which a given limb is in motion. Directly output by FLLIT. Values for each limb were pooled, giving six values per fly. |
| Stride Displacement | Straight-line distance between the posterior and anterior extreme positions of the limb in each stride. Directly output by FLLIT. Values for each limb were pooled, giving six values per fly. |
| Path travelled / displacement | Path Travelled (directly output by FLLIT) is the total actual distance covered by a limb during a stride; this is divided by Stride Displacement to compensate for differences in stride length. Values of 0 would indicate the limb moves in a perfectly straight line from posterior to anterior position, and larger values indicate greater deviation from this optimal path. Values for each limb were pooled, giving six values per fly. |
| Velocity | Path Travelled divided by Stride Duration (time between initiation and completion of a stride, directly output by FLLIT) to give the velocity of the limb. Values for each limb were pooled, giving six values per fly. |
| Domain overlap | Area of overlap ($\mu\text{m}^2$) between the domain (traced trajectories of each limb path) between each pair of limbs is directly output by FLLIT: % of flies in which there is at least one instance of non-zero overlap between any limb pair is presented. |
| Footprint regularity mid Legs | As in <sup>39</sup>. Average standard deviation (SD) of the posterior and anterior extreme absolute XY limb position: mean of the left and right mid-leg SDs. |
| Footprint regularity Hind legs | As in <sup>39</sup>. Average SD of the posterior and anterior extreme absolute XY limb position: mean of the left and right hind-leg SDs. |
| Domain length mid Legs | As in <sup>39</sup>. Maximum projected distance of the claw positions onto the major principal axis: mean of the left and right mid-legs. |
| Domain length hind Legs | As in <sup>39</sup>. Maximum projected distance of the claw positions onto the major principal axis: mean of the left and right hind-legs. |
| Stride length mid legs | As in <sup>39</sup>. As “Stride displacement”, mean of left and right mid-legs. |
| Stride length hind legs | As in <sup>39</sup>. As “Stride displacement”, mean of left and right hind-legs. |

### Leg twitch analysis

5 min videos of flies in horizontally-placed behavioural tubes were taken with an android phone camera. Leg twitches were manually identified as per the criteria described previously ^1^: briefly, “leg twitches” occurred only in a single limb, consisted of >1 repetitive movements of similar characteristics, did not involve grooming, and were not coincident with forward or backward locomotion of >1/2 body length. These leg twitches tended to occur in bouts; the number and mean duration of bouts per fly were quantified.

### Additional notes on DYSC-based thermogenetic control of SLO BK channel expression

The *dysc* locus encodes a scaffolding protein containing three PDZ domains ^27^. DYSC proteins bind to SLO BK channels and promote their functional expression in fly neurons in a post-translational manner, a process that likely requires the third C-terminal PDZ binding domain of DYSC ^27^. The expression of DYSC isoforms containing this third PDZ domain can be blocked via homozygosity for a P-element inserted between exons encoding the 2^nd^ and 3^rd^ PDZ domains (*dysc*^s168^) ^27^, the LOF allele we use in this study. We note that, in *dysc*^s168/s168^ homozygotes, shorter DYSC isoforms containing PDZ domains 1 and 2 are still transcribed, but SLO BK channel expression is nonetheless substantially reduced ^27^. Other LOF alleles of *dysc* are available – for example, the null allele *dysc*^c03838^ ^27^. However, homozygotes for *dysc* null alleles exhibit reduced viability (personal observations) and reduced peak movement ability ^27^. In contrast, the overall activity in *dysc*^s168^ homozygotes – as quantified by the number of infrared beam breaks over 24 h measured by the *Drosophila* Activity Monitor (DAM) system ^34^ – is slightly higher than isogenic control flies ^26^, illustrating that *dysc*^s168^ homozygotes (unlike *slo*^E366G/+^ BK channel GOF flies ^26^) do not exhibit severe motor defects. Hence, use of *dysc*^s168/s168^ homozygote flies as a genetic background suppressive for SLO BK channel expression enabled us to maintain the *slo*^E366G/+^ allele in a background with robust overall movement. This in turn allowed us to readily measure decreases in total movement caused by the re-expression of DYSC (and therefore SLO GOF BK channels harbouring the E366G mutation) during the key developmental window in which GOF BK channels act to perturb movement and limb kinematics.

### Statistical Analyses

Statistical analyses were performed using Graphpad Prism. Data sets were tested for normal distributions using the Shapiro-Wilk test. Normally distributed data sets were tested for statistical differences using unpaired t-tests with Welch’s correction for non-identical variance or one- or two-way ANOVA with Sidak’s multiple comparisons test; non-normally distributed data sets were tested using Mann-Whitney U-test or Kruskal-Wallis test with Dunn’s post-hoc test. Binary data were compared using chi-squared tests. Cohen’s effect size was calculated using the following formula: Cohen’s effect size *d* = (M_2_-M_1_)/SD_pooled_, where M_2_ and M_1_ are means of experimental vs control datasets, and SD_pooled_ = √((SD ^2^ + SD ^2^)/2), where SD_1_ + SD_2_ are the standard deviation of the experimental and control datasets. The coefficient of variation (CV) was calculated using the formula CV = SD/M, where SD is standard deviation of the data and M is the mean of the data.

