## Supplemental information for "BK channel gain-of-function disrupts limb control by suppressing neurotransmission during a critical developmental window"

**Supplementary Figures, Supplementary Figure Legends, and Supplementary References**

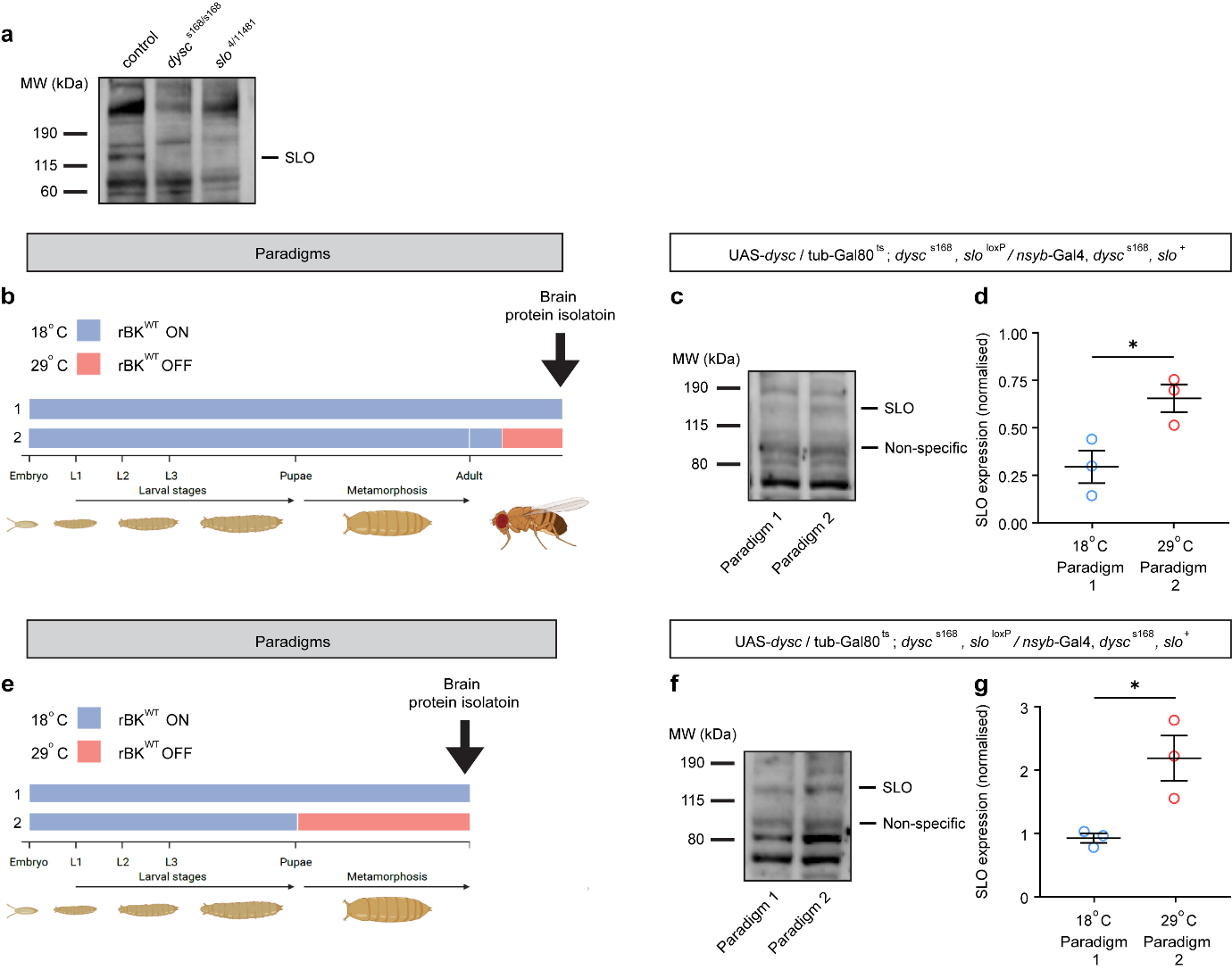

**Supplementary Fig. 1. Spatio-temporal control of SLO protein levels via dynamic regulation of DYSC expression.**

**(a)** Representative western blot showing SLO protein detected via an anti-hSlo1 antibody. A protein band was detected at the correct predicted molecular weight (125-134 kDa depending on alternative splice isoform) from control (iso31) brain tissue, and which was absent in slo null (*slo*^4^/*slo*^11481^) head tissue. The expression of SLO was also clearly reduced in head tissue from *dysc*^s168^ homozygotes.

**(b)** Temperature paradigm for inducing adult-stage rSLO^WT^ expression through temporal control of DYSC.

**(c-d)** Representative western blot (c) and quantification via densitometry of SLO expression (d) when GAL80^ts^-mediated suppression of neuronal DYSC expression is either active (18°C) or inactive (29°C) during days 3-5 of adulthood.

**(e)** Temperature paradigm for inducing pupal-stage rSLO^WT^ expression through temporal control of DYSC.

**(f-g)** Representative western blot (f) and quantification via densitometry of SLO expression (g) when GAL80^ts^-mediated suppression of neuronal DYSC expression is either active (18°C) or inactive (29°C) during pupariation. Brain samples were collected from pharate adult (late-stage) pupae subject to both temperature-paradigms.

Error bars: SEM. *p<0.05, student’s t-test. n = 3 independent protein samples derived from n = 10 brains for each genotype/temperature paradigm.

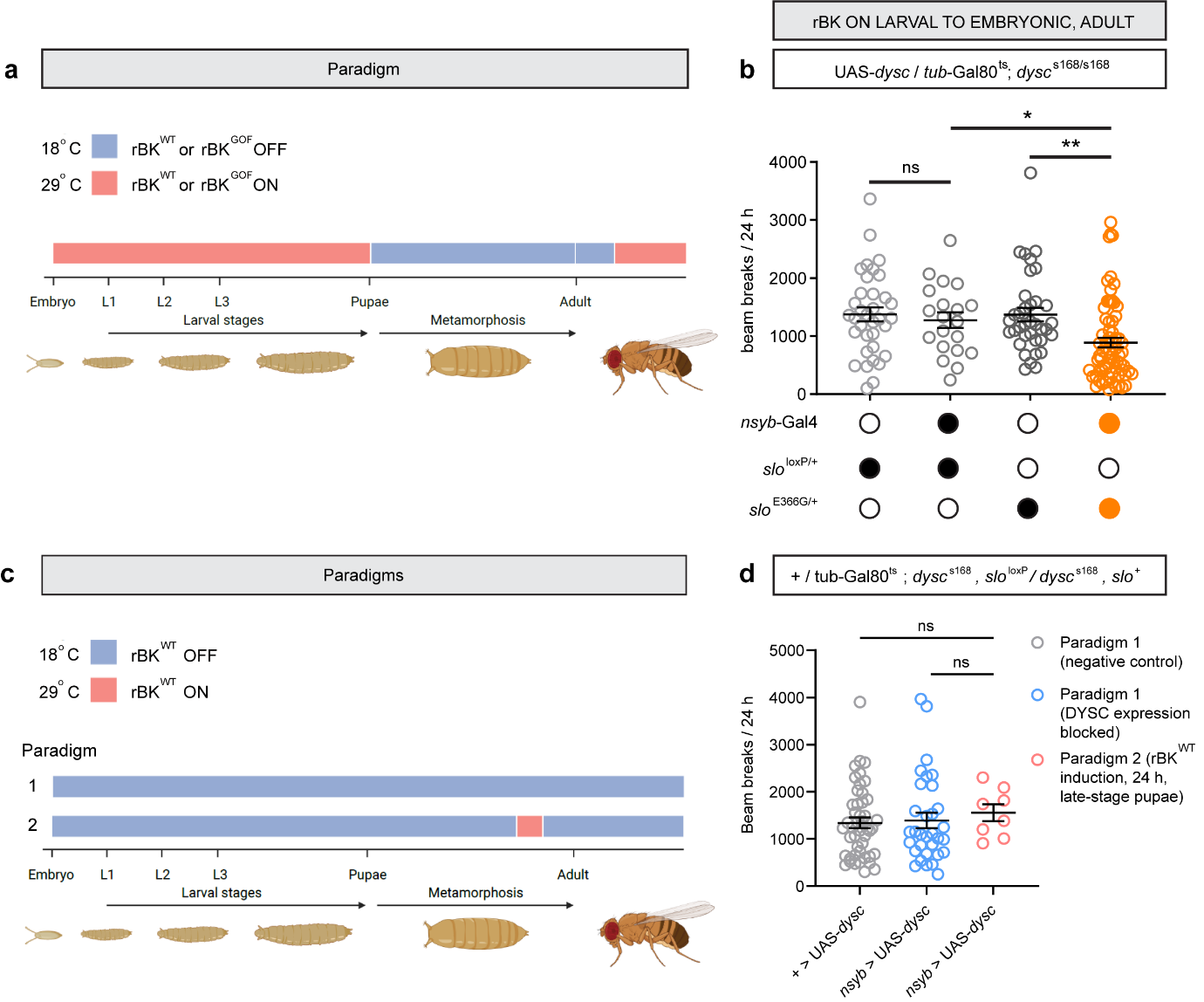

**Supplementary Fig. 2. Neuronal expression of GOF BK channels in all stages of the life cycle apart from the pupal stage causes a mild reduction in spontaneous locomotion**

**(a)** Schematic illustrating the stages of the *Drosophila* life cycle during which wild-type or GOF BK channels robustly expressed (29°C, red) or suppressed (18°C, blue). SLO BK channels were robustly expressed from the embryonic stage throughout the larval stage up until white pre-pupal formation; suppressed for the duration of the pupal stage; and robustly expressed again from 2 days post-eclosure for at least 3 days before experimentation, as well as throughout the duration of the experiment. Experiments on adult males took place 5-7 days post-eclosure.

**(b)** Total number of beam breaks in a 24 h period, recorded using the DAM system, in flies subject to the paradigm illustrated in (a). Control and genotypes are denoted in grey, experimental in orange. n values are (left to right): 35, 20, 36, and 70.

**(c)** Schematic illustrating the stage of the *Drosophila* life cycle during which wild-type BK channels are robustly expressed (29°C, red) or suppressed (18°C, blue). Experiments on adult males took place 5-7 days post-eclosure.

**(d)** Total number of beam breaks in a 24 h period, recorded using the DAM system, in flies subject to the paradigms illustrated in (c). n values are (left to right): 46, 32, 8.

Error bars: SEM. *p < 0.05, **p < 0.005, ns – p > 0.05, Kruskal-Wallis test with Dunn’s multiple comparisons.

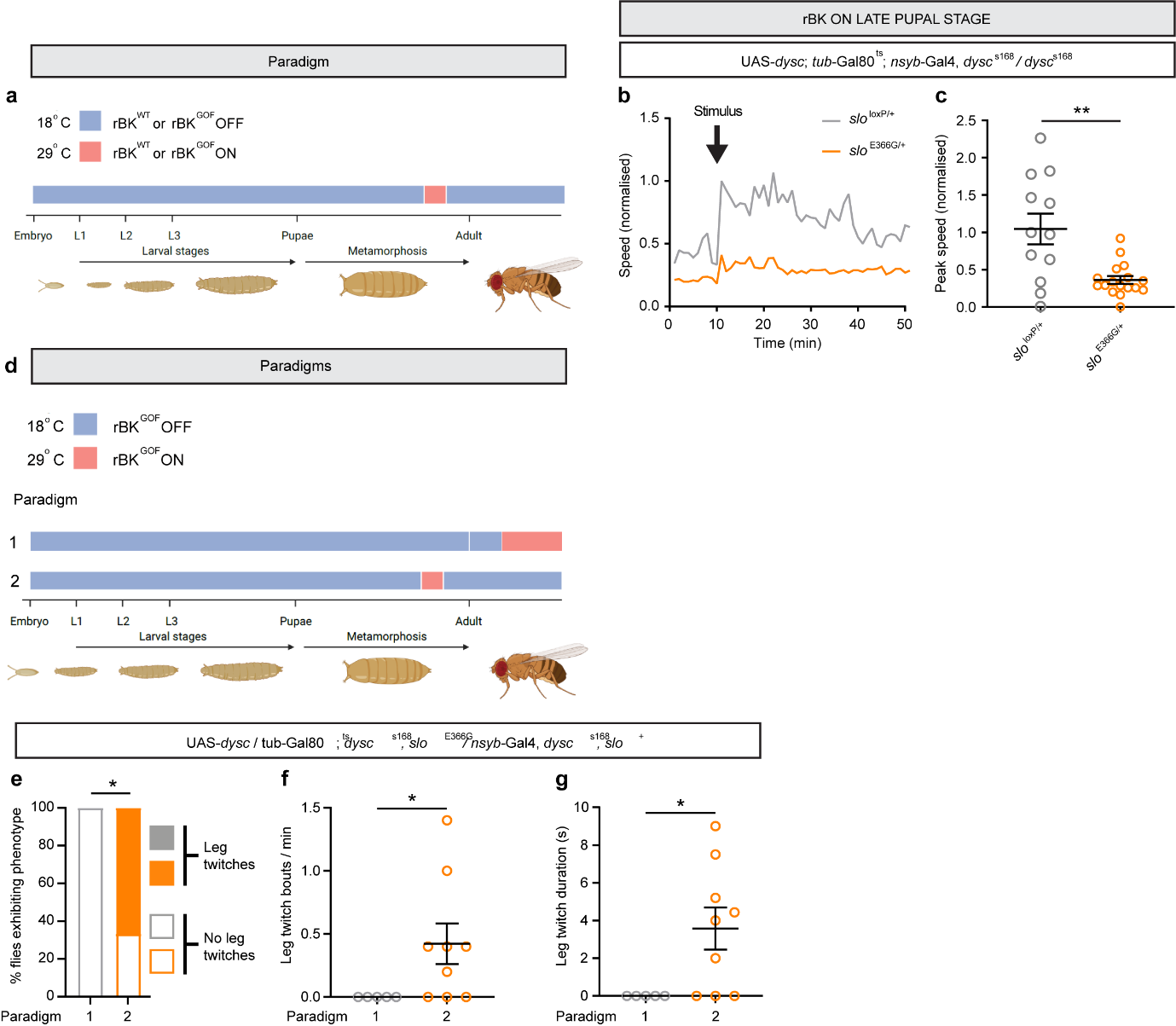

**Supplementary Fig. 3. Neuronal expression of GOF BK channels during a late developmental period reduces stimulus-dependent locomotion.**

**(a)** Schematic illustrating the stages of the *Drosophila* life cycle during which wild-type or GOF BK channels are robustly expressed (29°C, red) or suppressed (18°C, blue). Robust SLO expression is restricted to a 24 h period towards the end of the pupal stage, ending shortly before eclosure. Experiments on resulting adult male flies took place 5-7 days after eclosure.

**(b)** Traces showing average speed over time in 1 min bins (mm / s) for adult flies with robust neural expression of wild-type (n = 12) or GOF BK channels (n = 17) induced during late neurodevelopment. Mechanical stimulus is delivered at 10 mins.

**(c)** Quantification of speed following mechanical stimulus. Each data point represents average speed (mm / s) in 1 min bin immediately following stimulation for one fly. Data are presented normalised to the control mean. n = 12, 17.

**(d)** Schematic illustrating the stages of the *Drosophila* life cycle during which GOF BK channels are robustly expressed (29°C, red) or suppressed (18°C, blue). Experiments on resulting adult male flies took place 5-7 days after eclosure.

**(e)** % of adult flies with robust neural expression of GOF BK channels induced during adulthood (paradigm 1; n = 5) or late neurodevelopment (paradigm 2; n = 9) exhibiting at least one bout of leg twitches during a 5 min video.

**(f)** Number of leg twitches per minute in adult flies with robust neural expression of GOF BK channels induced during adulthood (paradigm 1; n = 5) or late neurodevelopment (paradigm 2; n = 9) during a 5 min video.

**(g)** Mean duration of leg twitch bouts in adult flies with robust neural expression of GOF BK channels induced during adulthood (paradigm 1; n = 5) or late neurodevelopment (paradigm 2; n = 9) during a 5 min video.

Error bars: SEM. *p< 0.05, **p< 0.005, Mann-Whitney U-test (c, f, g) or chi-squared test (e).

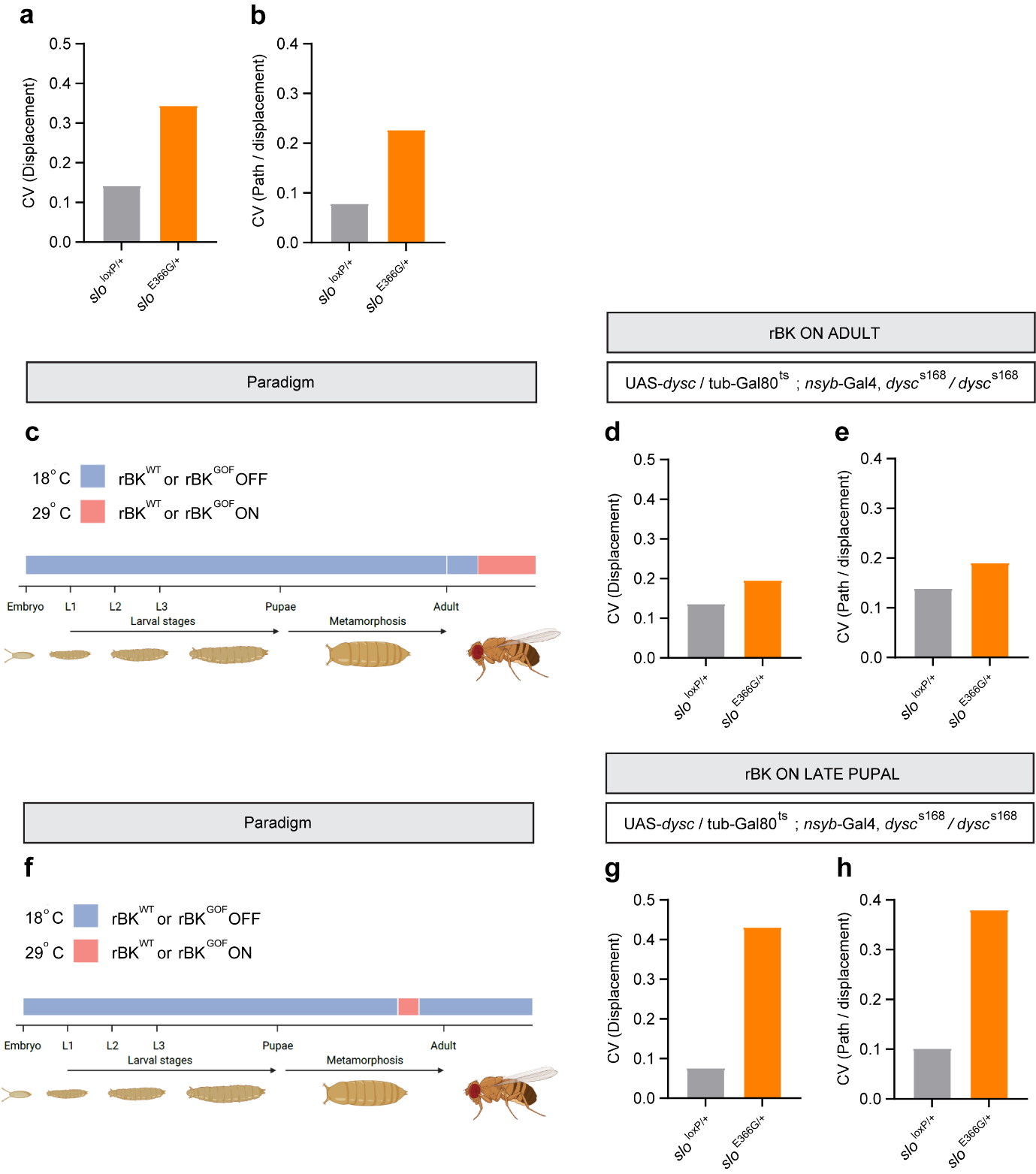

**Supplementary Fig. 4. Expression of GOF BK channels during neurodevelopment reduces the stereotypy of limb movements.**

**(a-b)** Coefficient of variance (standard deviation / mean) of FLITT-derived stride displacement comparing control (*slo*^loxP/+^) to BK GOF (*slo*^E366G/+^) flies. n = 66 limbs across 11 flies (*slo*^loxP/+^) and 60 limbs across 10 flies (*slo*^E366G/+^). Parameters are (a) Stride displacement and (b) Path travelled / stride displacement.

**(c)** Schematic illustrating the stages of the *Drosophila* life cycle during which wild-type or GOF BK channels are robustly expressed (29°C, red) or suppressed (18°C, blue). Experiments on resulting adult male flies took place 5-7 days after eclosure.

**(d-e)** Coefficient of variance of FLITT-derived stride displacement comparing flies with robust neural expression of wild-type (n = 54 limbs across 9 flies) or GOF BK channels (60 limbs across 10 flies) induced in mature adult neurons. Parameters are (d) Stride displacement and (e) Path travelled/ stride displacement.

**(f)** Schematic illustrating the stages of the *Drosophila* life cycle during which wild-type or GOF BK channels are robustly expressed 29°C, red) or suppressed (18°C, blue). Experiments on resulting adult male flies took place 5-7 days after eclosure.

(**g-h)** Coefficient of variance of FLITT-derived stride displacement comparing flies with robust neural expression of wild-type (n = 24 limbs across 4 flies) or GOF BK channels (42 limbs across 7 flies) induced during late neurodevelopment.

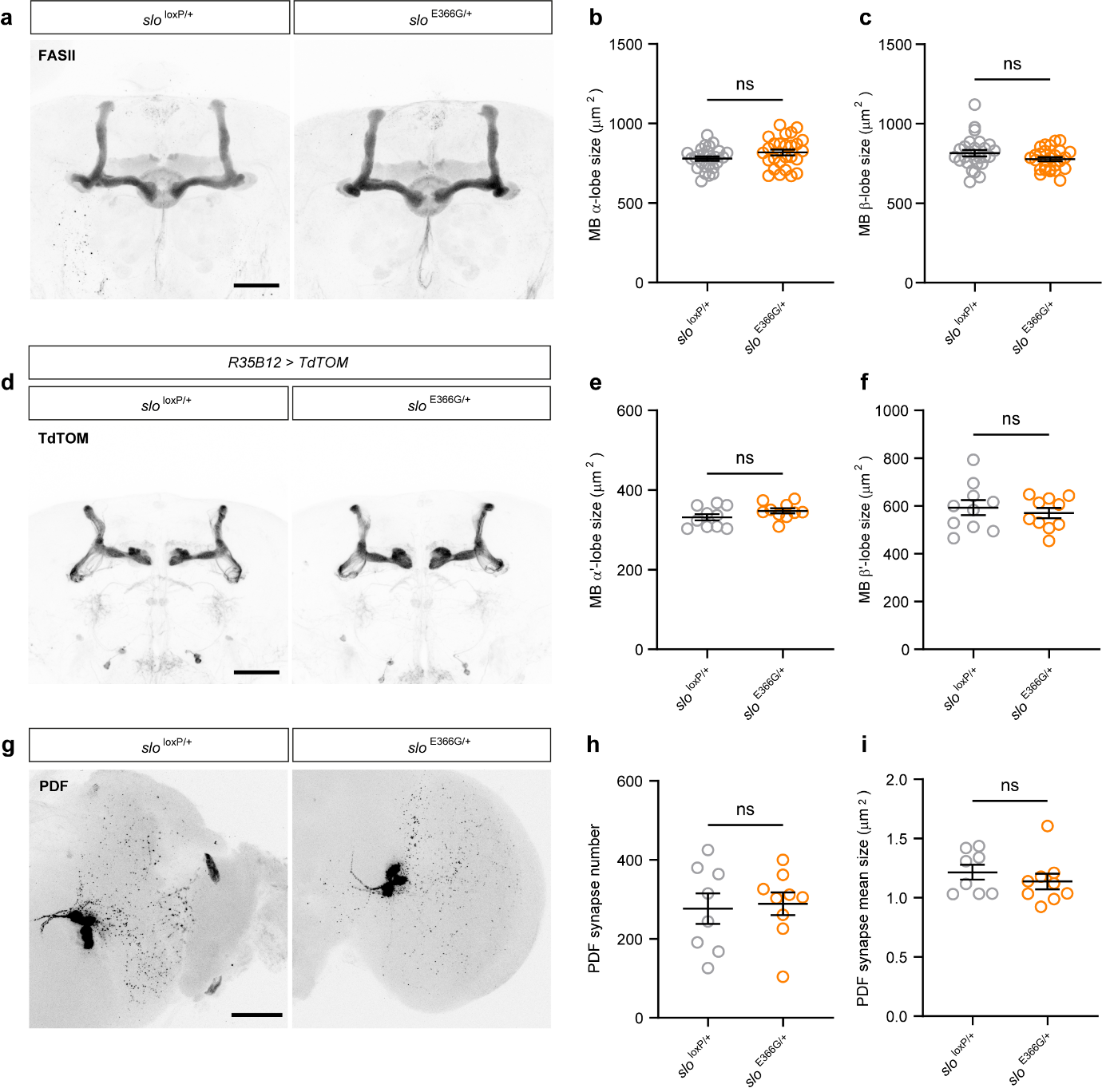

**Supplementary Fig. 5. Axonal outgrowth in mushroom body neurons, and synaptic development in PDF-positive l-LN_v_ neurons, are unaltered in flies expressing GOF BK channels.**

**(a)** Representative images of FASII immuno-staining in control (*slo*^loxP/+^) and BK channel GOF (*slo*^E366G/+^) adult brains. Both vertical (α) and horizontal (β) lobes of the mushroom bodies are strongly labelled by FASII antibodies. Scale bar: 50 µm.

**(b-c)** Quantification of α-lobe (b) and β-lobe (c) area in *slo*^loxP/+^ and *slo*^E366G/+^ adult brains. n = 26 lobes from n = 13 flies for both genotypes.

**(d)** Representative images of TdTomato (TdTom) driven by the *R35B12*-Gal4 driver, which strongly labels the mushroom body α’- and β’-neurons ^1^, in *slo*^loxP/+^ and *slo*^E366G/+^ adult brains. Scale bar: 50 µm.

**(e-f)** Quantification of vertical α’-lobe (E) and horizontal β’-lobe (F) area in *slo*^loxP/+^ and *slo*^E366G/+^ adult brains. n = 10 lobes from n = 5 flies for both genotypes.

**(g)** Representative images of PDF immuno-staining in *slo*^loxP/+^ and *slo*^E366G/+^ adult optic lobes. Strongly stained cell bodies of PDF-positive s-LN_v_ and l-LN_v_ neurons are visible. PDF-positive synaptic puncta of l-LN_v_ neurons are decorated throughout the optic lobes. Scale bar: 50 µm.

**(h-i)** Number (h) and mean area per optic lobe (i) of PDF-positive synaptic puncta in *slo*^loxP/+^ and *slo*^E366G/+^ adult optic lobes. *slo*^loxP/+^: n = 8 optic lobes from n = 4 flies; *slo*^E366G/+^: n = 9 optic lobes from n = 5 flies.

Error bars: SEM. ns – p > 0.05, t-test with Welch’s correction (b, c, e, f, h) or Mann-Whitney U-test (i).

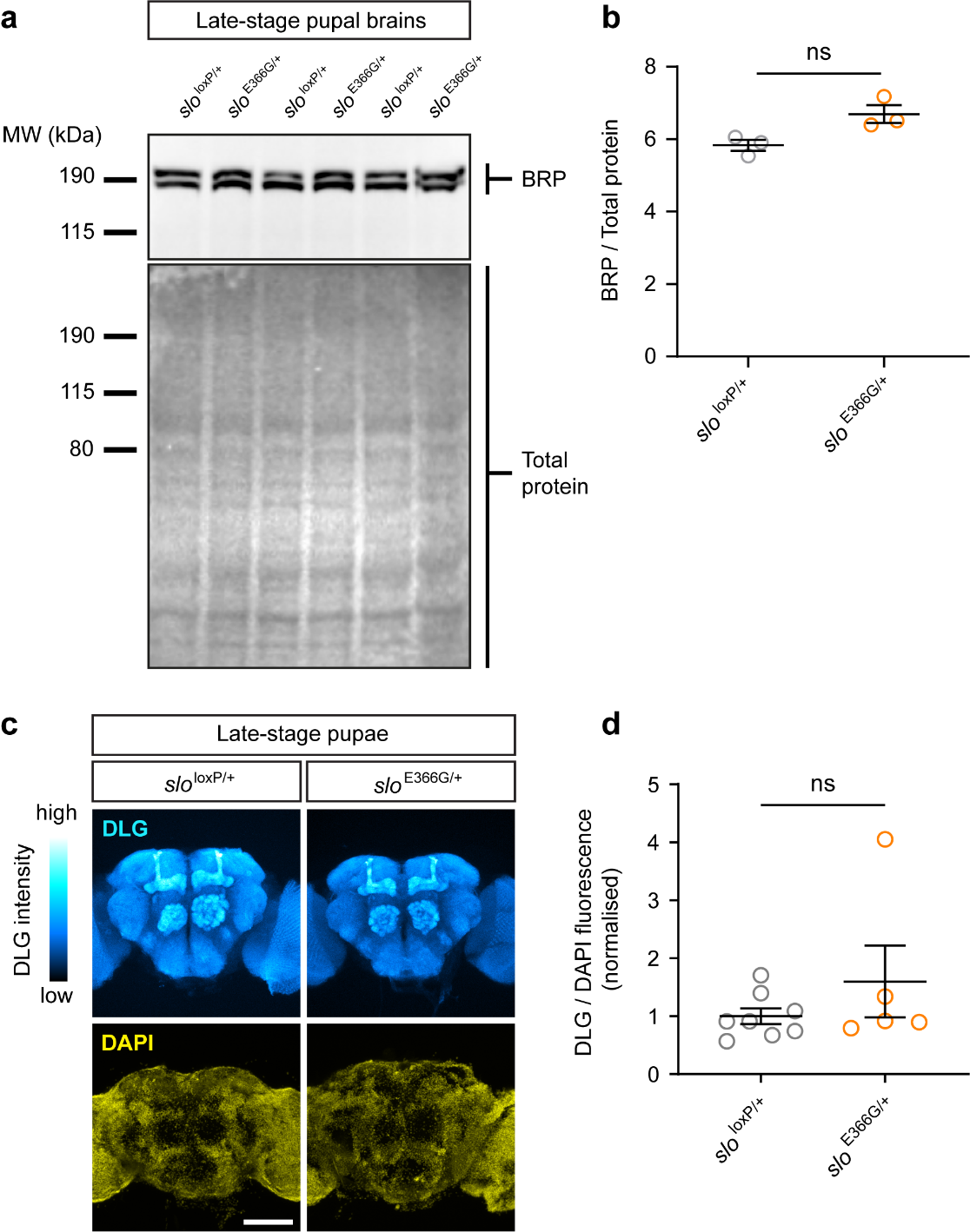

**Supplementary Fig. 6. Bruchpilot (BRP) protein levels and immuno-fluorescence of Discs Large (DLG) are unaltered by BK channel GOF.**

**(a)** Western blot data comparing BRP protein levels in 3 biological replicates of control (*slo*^loxP/+^) and BK channel GOF (*slo*^E366G/+^) late-stage pupal brains (n = 10 per sample).

**(b)** Quantified BRP levels, normalised to total protein, in *slo*^loxP/+^ to *slo*^E366G/+^ late-stage pupal brains. Overall BRP levels were marginally, but significantly, enhanced by BK channel GOF.

**(c)** Representative images of control (*slo*^loxP/+^) and BK channel GOF (*slo*^E366G/+^) late-stage pupal brains immuno-stained with anti-DLG and counterstained with DAPI.

**(d)** Quantified whole-brain DLG immunofluorescence, normalised to DAPI, in control (*slo*^loxP/+^; n = 8) and BK channel GOF (*slo*^E366G/+^; n = 5) late-stage pupal brains. Data are shown normalised to the mean of *slo*^loxP/+^ controls.

Error bars: SEM. * p < 0.05, ns – p > 0.05, student’s t-test (b) or Mann-Whitney U-test (d).

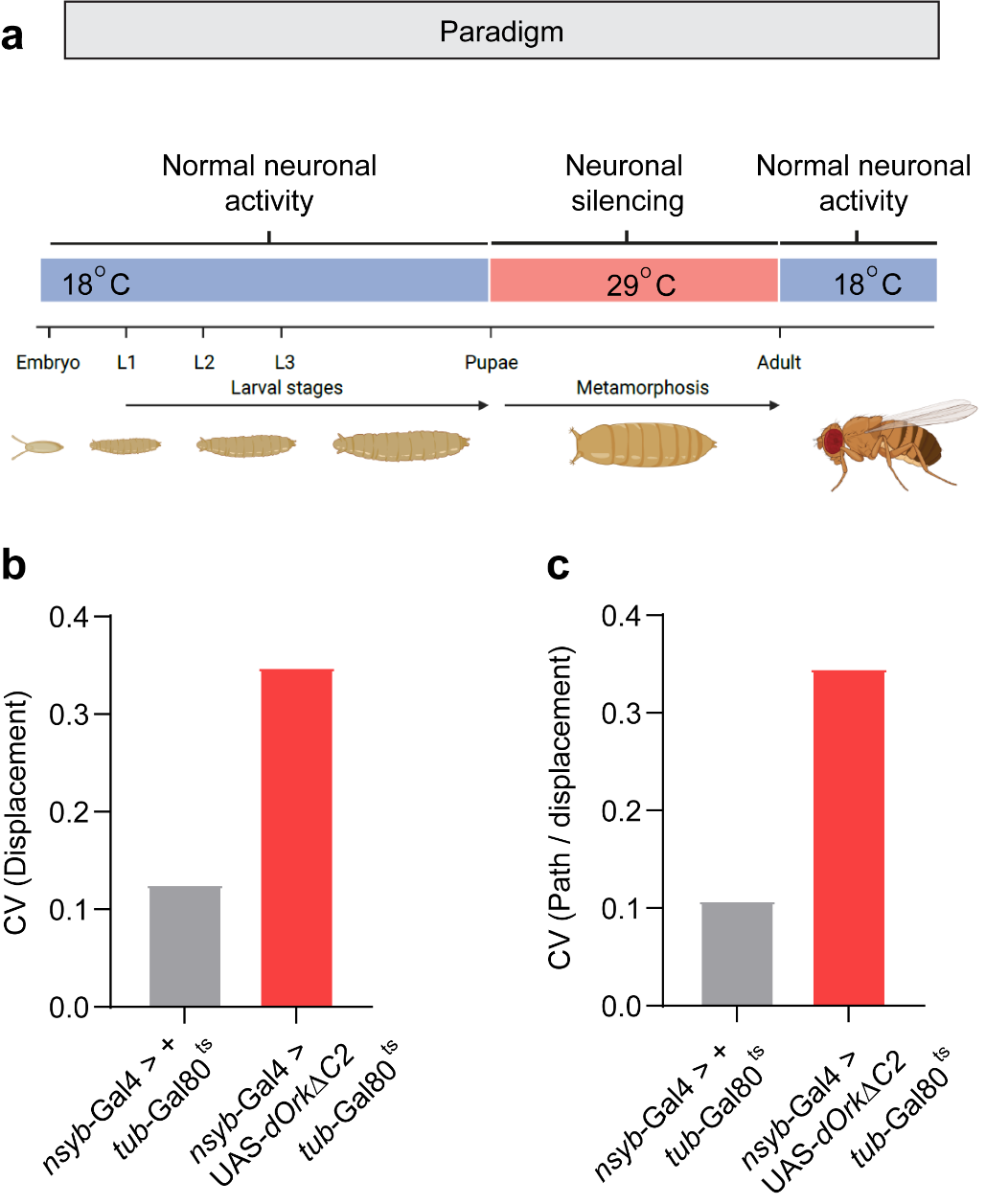

**Supplementary Fig. 7. Neuronal silencing during development reduces the stereotypy of limb movements.**

**(a)** Schematic illustrating the stages of the *Drosophila* life cycle during which dOrkΔC2 open rectifying channels (which suppress neuronal excitability) were robustly expressed (29°C, red) or suppressed (18°C, blue). Experiments on resulting adult male flies took place 5-7 days after eclosure.

**(b, c)** Coefficient of variance (standard deviation / mean) of FLITT-derived stride displacement comparing control adults (*nsyb-*Gal4 */ +, tub-*Gal80^ts^ / +) and adults subject to pupal-stage neuronal silencing (*nsyb-*Gal4 */ +, tub-*Gal80^ts^ / UAS-*dOrkΔC2*) flies. n = 42 limbs across 7 flies for both genotypes. Parameters are (b) Stride displacement and (c) Path travelled / stride displacement.

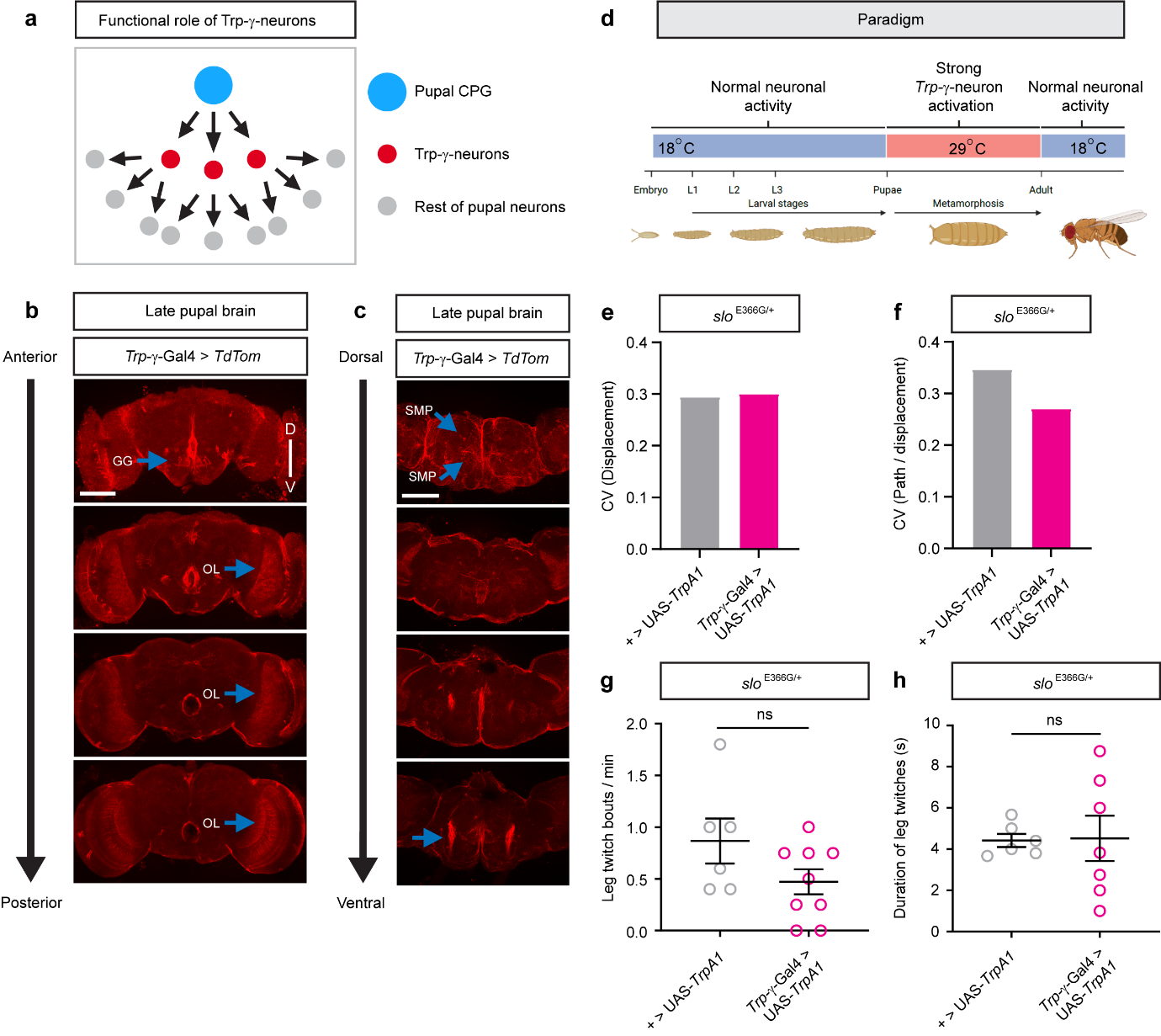

**Supplementary Fig. 8. Trp-γ-neurons in the pupal *Drosophila* brain.**

**(a)** Proposed functional role of pupal Trp-γ-neurons (based on ^2^). Trp-γ-neurons may act as components of a relay network that transmits excitatory signals from an undefined central pattern generator (CPG) to the remainder of the pupal nervous system, thus promoting stimulus-independent neural activity during development of the adult *Drosophila* nervous system.

**(b-c)** Anterior to posterior (b) and dorso-ventral (c) confocal stacks (20-30 µm in depth) illustrating the location and projection patterns of late-stage pupal Trp-γ-neurons, labelled using the *Trp*-γ-Gal4 reporter to drive a membrane tagged fluorescent protein (UAS-CD4::TdTomato; UAS-*TdTom*). Dorso-ventral (D-V) and anterior to posterior (A-P) axes are also shown in (b) and (c) respectively. Cell bodies of late-stage pupal Trp-γ-neurons can be observed in several neuropil domains, including the gnathal ganglia (GG), lamina and medulla neuropils of the optic lobe (OL), and dorsal regions of the superior medial protocerebrum (SMP). Dense projections arising closely to the GG and projecting to the center of the pupal brain can be clearly observed (bottom arrow, c).

**(d)** Schematic illustrating the stages of the *Drosophila* life cycle during which *Trp-γ-*neurons were subject to strong excitation via TrpA1 activation (29°C, red) or were not subject to manipulation (18°C, blue). Experiments on resulting adult male flies took place 5-7 days after eclosure.

**(e, f)** Coefficient of variance (standard deviation / mean) of FLITT-derived stride displacement (E) or the non-linearity of movement (F) in flies expressing GOF BK channels (*slo*^E366G/+^) that either were not subjected (UAS*-TrpA1* / +) or were subjected (*Trp-γ-*Gal4 *>* UAS*-TrpA1*) to pupal-stage excitation of Trp-γ*-*neurons flies. n = 78 limbs across 13 flies for both genotypes.

**(g)** Number of leg twitches per minute during a 5 min video in adult flies that either were not subjected (UAS*-TrpA1* / +; n = 6) or were subjected (*Trp-γ-*Gal4 *>* UAS*-TrpA1*; n = 9) to pupal-stage excitation of Trp-γ-neurons.

**(h)** Mean duration of leg twitch bouts during a 5 min video in adult flies that either were not subjected (UAS*-TrpA1* / +; n = 6) or were subjected (*Trp-γ-*Gal4 *>* UAS*-TrpA1*; n = 7) to pupal-stage excitation of Trp-γ-neurons.
